# Age determines NK cell fate and tissue compartmentalization to CMV infection

**DOI:** 10.64898/2026.03.16.712099

**Authors:** Isaac J. Jensen, Benjamin J. Redenti, Steven B. Wells, Joshua I. Gray, Daniel P. Caron, Donna L. Farber

## Abstract

Cytomegalovirus (CMV), a ubiquitous herpesvirus, establishes persistent infection that is controlled by both NK and CD8 T cells. While immunity to CMV is primarily studied in adult mice and humans, most individuals acquire CMV during the early years of life and virus-specific NK and T cell responses during this vulnerable life stage are understudied. Here, we show distinct responses by infant and adult NK cells to CMV infection in both mice and humans resulting from cell-intrinsic features and host T cell responses. Infant mice sustained higher viral loads compared to adults following MCMV infection and exhibited non-redundant requirements for NK cells. Infant MCMV-reactive (Ly49H^+^) NK cells preferentially expanded adaptive-like subsets which were maintained in tissues and exhibited a distinct transcriptional profile relative to adult NK cells. This biased differentiation of adaptive-like NK cells was altered over age and controlled in part, by competition with memory T cells. We demonstrate similar dynamics with human NK cells; distinct adaptive-profiles for HCMV-reactive NK cells in early life and childhood that change over age and are inversely associated with anti-viral T cell responses. Together, our results reveal that NK cells develop adaptive-like responses and seed tissues in early life to provide protective memory when T cell immunity is limited.

**ONE-SENTENCE SUMMARY:** Memory NK cell generation to cytomegalovirus infection is regulated by age and T cell immunity

## INTRODUCTION

Human cytomegalovirus (HCMV) infection is globally endemic with a seroprevalence ranging from 60-90% worldwide (*1*). In adults, acute and persistent CMV infection is often asymptomatic and can be controlled by a healthy immune system. During fetal development however, HCMV infection is the most common congenital infection and can result in devastating outcomes such as infant deafness (*2–5*). During infancy, HCMV infection is ubiquitous, infecting 25% of children in the first year (accounting for nearly half of all HCMV infections) and the incidence of childhood HCMV infections is increasing in the USA (*6*, *7*). Despite the prevalence of early life infection, the immune response to CMV is mostly studied in adult mice and humans. The mechanisms by which the infant immune system responds to CMV infection are unresolved and critical for developing age-appropriate strategies for protection.

Persistent CMV infection is kept in check by ongoing immune effector responses. Humans born lacking either Natural Killer (NK) or CD8^+^ T cells develop severe complications following exposure to HCMV and other Herpes viruses (*8–10*). Both NK cells and CD8^+^ T cells robustly target virally infected cells for cytolysis (*11*, *12*). However, specific responses by NK or CD8^+^ T cells are usually studied separately and the coordinate role of each subset in viral protection, differentiation and memory formation in the context of early life infections remain unexplored.

NK cells are considered part of the innate immune system because they mediate early responses to pathogens through rapid and direct cytolytic activity and broad recognition of histocompatibility molecules (*13–16*). In the case of CMV infection however, NK cells can exhibit adaptive-like features by expression of receptors that respond specifically to CMV in mice and humans (*17–20*). Namely, the killer lectin-like receptor (KLR) Ly49H (*Klra8*) confers specificity for MCMV (*19*) thereby limiting infection morbidity (*14*, *20–22*). Following MCMV infection, Ly49H^+^ NK cells persist as a phenotypically distinct subset exhibiting a KLRG1^+^ CD62L^-^ phenotype and exhibiting certain features of adaptive-like memory cells in their ability to mediate enhanced responses and protection against MCMV and their maintenance in tissue reservoirs, such as the liver (*20*, *21*, *23*, *24*). In humans, NKG2C expression marks HCMV-reactive NK cells, which have increased cytotoxicity and are enriched in the blood of HCMV seropositive individuals (*17*, *25*, *26*)— similarly suggesting a type of NK cell memory. However, a consensus signature for human adaptive-like NK cells in tissues has not been elucidated and their dynamics across age remains unexplored.

CD8^+^ T cell responses to CMV infection peak after the NK cell responses to mediate viral clearance and subsequently develop long-lived, heterogeneous memory populations. In mice, MCMV-specific memory CD8^+^ T cells can persist in multiple tissues (including infection sites) as conventional resting memory subsets specific for epitopes expressed during acute infection, or “inflationary memory” populations specific for persisting viral epitopes(*27–29*). In humans, HCMV-reactive memory T cells are detected in increasing frequencies over age and exhibit a predominant terminal effector-like or T_EMRA_ profile as measured in blood and other sites like the bone marrow (*30*, *31*). T cell responses in early life CMV infection have not been comprehensively studied, though in acute infections (influenza, listeria), mouse infant T cells are biased toward effector responses with limited capacity for establishing memory compared to adult T cells (*32–36*). Whether adaptive-like NK cells may compensate for the lack of T cell memory in early life CMV infections has not been assessed.

In this study, we investigated the NK cell response to CMV infection in early life relative to adults including its role in protection, propensity to generate adaptive-like responses, and interactions with T cell immunity using the MCMV infection model and novel human tissue samples. In mice, NK cells provided early protection to disseminated MCMV infection during infancy while T cells control infection at later times. Infant MCMV-reactive (Ly49H^+^) NK cells preferentially expanded adaptive-like subsets which were maintained in tissues and exhibited a distinct transcriptional profile relative to adult NK cells. With age, the propensity to maintain adaptive-like NK cells diminished coincident with the increased capacity of T cells to form durable memory responses. In humans, we identified a transcriptional signature for adaptive-like NK cells that predominates in tissues of HCMV-individuals. Human adaptive-like NK cells in early life resembled infant MCMV-specific NK cells, and over age accumulate CD57 expression, a marker of repeated stimulation. Frequencies of HCMV-reactive NK cell memory are inversely proportional to that of HCMV-expanded CD8^+^ T_EMRA_ cells. Together, our results reveal that CMV infection at early ages establishes long-lived NK cell memory in tissues and provides crucial protection when T cell memory is lacking.

## RESULTS

### Infant mice require both NK and CD8^+^ T cells to limit infection morbidity

To study the dynamics of MCMV infection during infancy, we infected infant (10 days post-natal [DPN]) and adult (8-week-old) mice intraperitoneally (i.p.) with MCMV and assessed viral load in blood and tissues between days 3-15 post-infection (**Fig. 1A**). Infant mice were infected at 10 DPN, this age more accurately reflects the human infant immune system than neonatal mice which are lymphopenic at birth (*33*). Tissues examined included sites of early viral replication (liver [LVR], lung [LNG]), sites of virus persistence (salivary gland [SVG], bone marrow [BOM]), lymphoid organs for immune surveillance of infected tissues (spleen [SPL], cervical lymph node [CLN]), and plasma, indicating hematogenous spread of the virus (*12*, *37*). Infant mice had consistently higher MCMV viral burdens across all sites starting at early times post-infection; virus was mostly cleared by 15 days post-infection (DPI) except in the SVG, a known site of viral persistence (*38*), where viral load increased from 3-15 DPI (**Fig. 1A**). These data demonstrate increased viral burden for MCMV infection in infants as in a previous study (*39*), and reveals that the virus disseminates across multiple sites including but not limited to known reservoirs for viral persistence.

**Figure 1.**
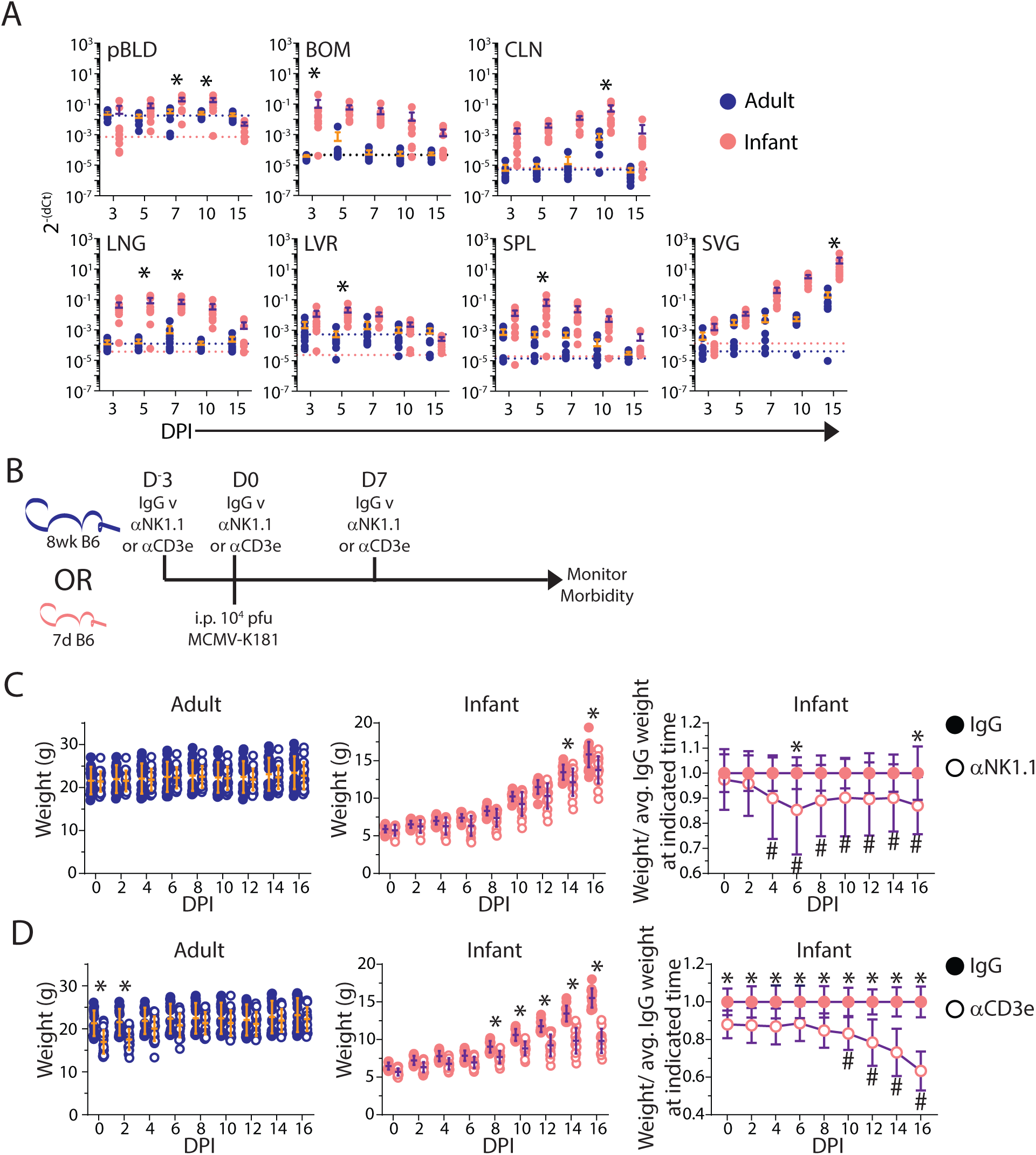
Infant mice have greater systemic viral burden and require both NK and T cells to limit morbidity. (A)Viral burden across tissues including blood plasma (pBLD), bone marrow (BOM), cervical lymph node (CLN), lung (LNG), liver (LVR), spleen (SPL), and salivary gland (SVG) at indicated days post infection (DPI). n = 7-10 mice per group. Blue and pink dotted lines represent limit of quantification for adult and infant tissues, respectively. (B) Experimental design for panels C and D. Weight of MCMV infected adult (left) and infant (middle) mice treated with control IgG (filled circle) or depleting antibody (open circle) for (C) NK cells [αNK1.1] or (D) T cells [αCD3e]. (C right, D right) Infant weights normalized to average IgG weight at indicated time. n = 20 mice per group. All error bars are mean ± SD. * p-value < 0.05 between groups comparison at indicated time-point; # p-value < 0.05 compared to D0 as assessed by two-way ANOVA with Bonferroni multiple comparisons test. All data are combined from N = 2 independent experimental replicates.

Both NK cells and T cells are potent cytolytic effectors which each play important roles in controlling MCMV infection (*12*, *40*). To dissect their relative contributions for immune-mediated protection to MCMV infection during infancy relative to adults, we performed antibody-mediated depletion of NK or T cells, beginning 3 days prior to infection and up to16 DPI when viral loads have diminished in most sites (**Fig. 1B**). Adult mice did not exhibit weight loss morbidity to MCMV infection when either NK cells or T cells were depleted, maintaining normal weights throughout the infection course (**Fig. 1C**, left), consistent with redundancy in NK and CD8 T cells in immune-mediated protection (*41*). By contrast, depletion of either NK cells or T cells in infant mice resulted in significantly enhanced morbidity relative to control-treated mice starting at early times post-infection (4 DPI) for NK cell depletion and later times post-infection (10 DPI) for T cell depletion (**Fig. 1C, D**). Morbidity for NK or T cell depletion persisted throughout the course of infection reaching a nadir at 6 DPI for NK cell depletion and 16 DPI for T cell depletion (**Fig. 1C,D**). These results indicate that both NK cells and T cells play critical, non-redundant roles in immune responses to MCMV infection during early life; NK cells provide essential protection during the first week when T cells are dispensable, but T cells are required for maintaining infection control at later times.

### Kinetics of the adaptive NK cell responses to MCMV across tissues

Given their critical role in mediating protection to MCMV in early life, we investigated the kinetics of NK cell development in infant mice at early post-natal ages and their responses to MCMV infection relative to adult mice. At baseline in uninfected mice, the frequency and number of total NK cells and MCMV-reactive (Ly49H^+^) NK cells was low and negligible, respectively, starting at 10 days post-natal (DPN), though both populations increased substantially between 10 and 40 DPN across blood and all tissues examined, and remained stable thereafter (up to 120 days) (**fig. S2,** see **fig. S1A** for gating strategy). In response to MCMV infection, infant mice showed increased frequency and numbers of Ly49H^+^ NK cells in the blood, spleen, and salivary gland (SVG) that exceeded those in uninfected mice and exhibited site specific dynamics (peak increases at 15 DPI in blood and spleen and progressive increases from 40-60 DPI in SVG) (**Fig. 2A**). By contrast, in adults, the frequency of Ly49H^+^ NK cells was already high (>50%) at baseline and increased slightly at 15 DPI, returning to baseline thereafter in the blood and spleen, with increased populations in the SVG (**Fig. 2B**). Together, these results show dynamic production and seeding of NK cells in multiple sites during the early weeks of life while adults maintain stable numbers across sites.

**Figure 2.**
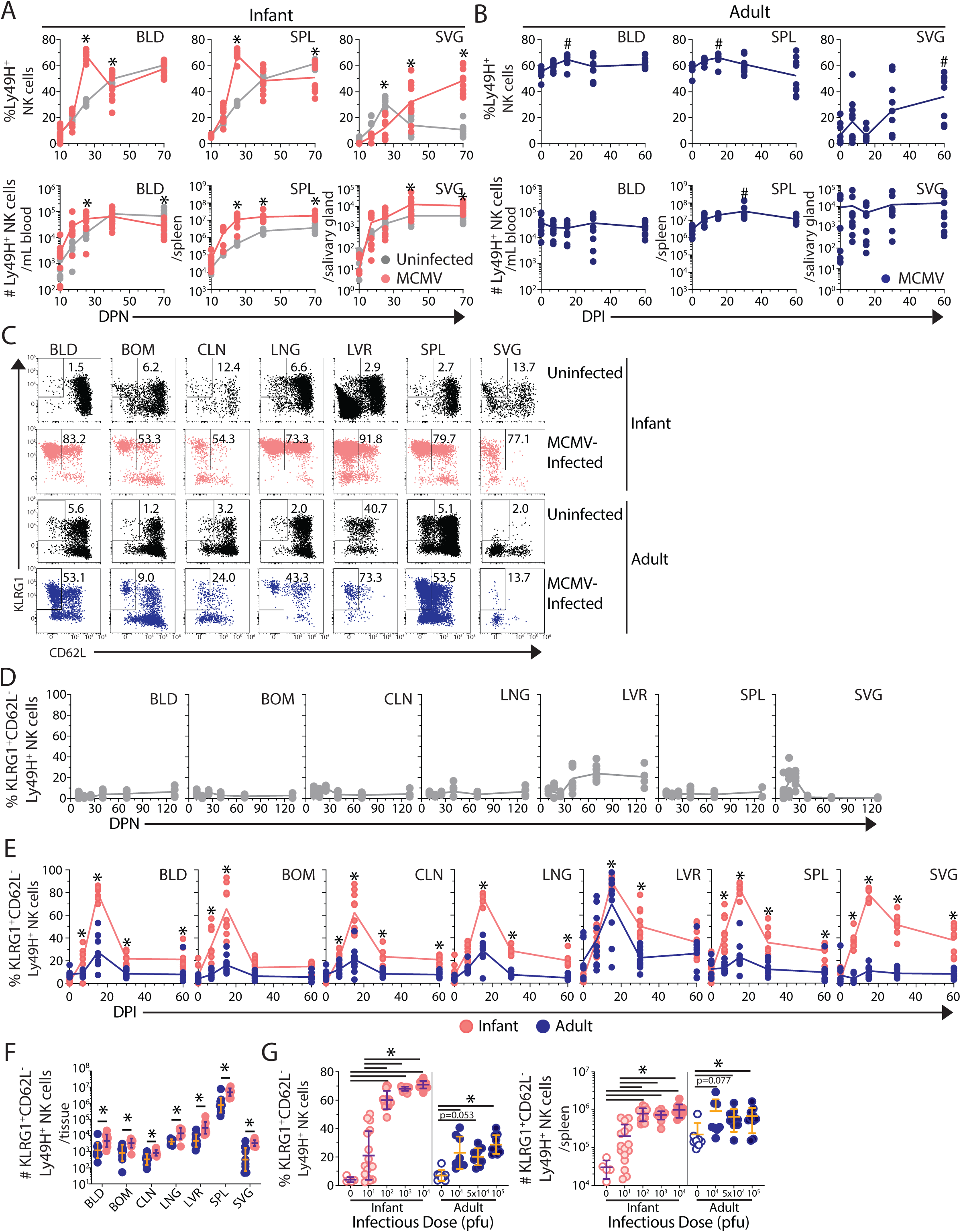
Dynamics of the NK cell response to MCMV across tissue. (A) Frequency (top) and number (bottom) of Ly49H^+^ NK cells in infection naive (grey) and MCMV-infected infant (pink) tissues at indicated post-natal days (DPN). *p-value < 0.05 as assessed by two-way ANOVA with Bonferroni multiple comparisons test. n = 8-10 mice per time-point per group. (B) Frequency and number of Ly49H^+^ NK cells in MCMV-infected adult (blue) tissues at indicated days post-infection (DPI). ^#^p-value < 0.05 as assessed by one-way ANOVA with Bonferroni multiple comparisons test relative to 0 DPI. n = 10 mice per time-point per group. (C) Representative gating of adaptive-like (KLRG1^+^ CD62L^-^) Ly49H^+^ NK cells from uninfected and MCMV-infected adult and infant tissues (15 DPI). (D) Frequency of adaptive-like Ly49H^+^ NK cells in indicated tissues from 10-130 DPN. n= 4-8 mice per timepoint. (E) Frequency of adaptive-like Ly49H^+^ NK cells in adult (blue) and infant (pink) tissues at indicated DPI. n = 10 mice per time-point per group. *p-value < 0.05 as assessed by two-way ANOVA with Bonferroni multiple comparisons test. (F) Number of adaptive-like Ly49H^+^ NK cells in adult (blue) and infant (pink) tissues at 60 DPI. n= 10 mice per group. *q-value <0.05 as assessed by corrected multiple t-statistic (false discovery rate [FDR]<5%) (G) Frequency (left) and number (right) of adaptive NK cells in spleen of infant (pink) and adult (blue) at 15 DPI with indicated infectious dose. n=5-16 mice per group. * p-value < 0.05 as assessed by separate one-way ANOVAs with Bonferroni multiple comparisons test. All data are combined from N = 2 independent experimental replicates. Tissue Abbreviations: blood (BLD), bone marrow (BOM), cervical lymph node (CLN), lung (LNG), liver (LVR), spleen (SPL), and salivary gland (SVG).

We then assessed whether MCMV-reactive (Ly49H^+^) exhibited an adaptive-like profile defined by KLRG1^+^ CD62L^-^ phenotypes (*23*) during development and following MCMV infection. In uninfected mice there were scant adaptive-like Ly49H^+^ NK cells across sites and ages, with a small proportion of KLRG1^+^ CD62L^-^ Ly49H^+^ NK cells in the liver starting at 40 DPN (**Fig. 2C,D**), consistent with the liver being a reservoir for adaptive-like or memory NK cells (*24*, *26*, *42*). Following MCMV infection of infant mice, there was a striking expansion of KLRG1^+^ CD62L^-^ Ly49H^+^ NK cells across all sites that peaked at 15 DPI and greatly exceeded their expansion following MCMV infection of adult mice, with the exception of the liver where adaptive-like NK cells were similarly expanded at 15 DPI in infant and adult mice (**Fig. 2C,E**). This enhanced frequency and numbers of adaptive-like NK cells persisted up to 60 DPI in BLD, SPL, LNG, CLN, and SVG in mice infected as infants compared to mice infected as adults (**Fig. 2E,F**). These results show that early life MCMV infection optimally mobilizes NK cells for robust virus-driven expansion and establishment of NK memory in multiple tissue reservoirs.

We investigated whether differences in viral burden drive the infant-specific expansion of adaptive-like NK cells by infecting infant and adult mice with graded viral doses (decreased for infants and increased for adults relative to the standard 10^4^ pfu dose used above) and assessing the NK response in the spleen at 15 DPI (peak expansion). However, infant mice infected with lower viral doses (<10^4^ pfu) still exhibited robust expansion and predominance of adaptive-like Ly49H^+^ NK cells, except for variability at the lowest dose (**Fig. 2G**). Similarly, increasing MCMV doses (>10^4^ pfu) in adult mice did not increase the frequency of adaptive-like NK cells (**Fig. 2G**). Overall, the differences between the infant and adult NK cell response were maintained across a 1000-fold variation in infectious dose (10^2^-infant, 10^5^-adult) exceeding the previously measured difference in viral burden **(Fig. 1A**). These results indicate that the propensity for adaptive-like NK cells expansion during infancy was intrinsic to the host.

### Distinct transcriptional profiles for infant and adult NK cells responding to MCMV

To assess potential mechanisms for the age-dependent NK cell responses to MCMV infection, we performed cellular indexing of transcriptome and epitopes (CITE-seq) of adult and infant NK cells prior to (0 DPI) and following (7, 15 DPI) infection, representing the acute and peak NK cell response to MCMV in infants **(Fig. 2**). Visualizing the dataset by UMAP embedding revealed variations based on infant and adult mice and time post-infection (**Fig. 3A**). There was also concordance between gene and protein expression for markers delineating adaptive-like NK cells (**Fig. 3B**). Differential expression analysis of adult and infant NK cells at each timepoint revealed distinct transcriptional profiles both prior to and following MCMV infection (**Fig. 3C** and **Table S1**). At baseline (0 DPI) infant NK cells had greater expression of genes associated with NK cell precursors (*Tcf7*, *Il7r*, *Rora*)(*43*) and proliferation (*Mki67*, *Tubb5*, *Kif1c*, *Kif2c*, *etc*) and lower expression of genes for effector function (*Ccl3*, *Ccl4*, *Ccl5*).

**Figure 3.**
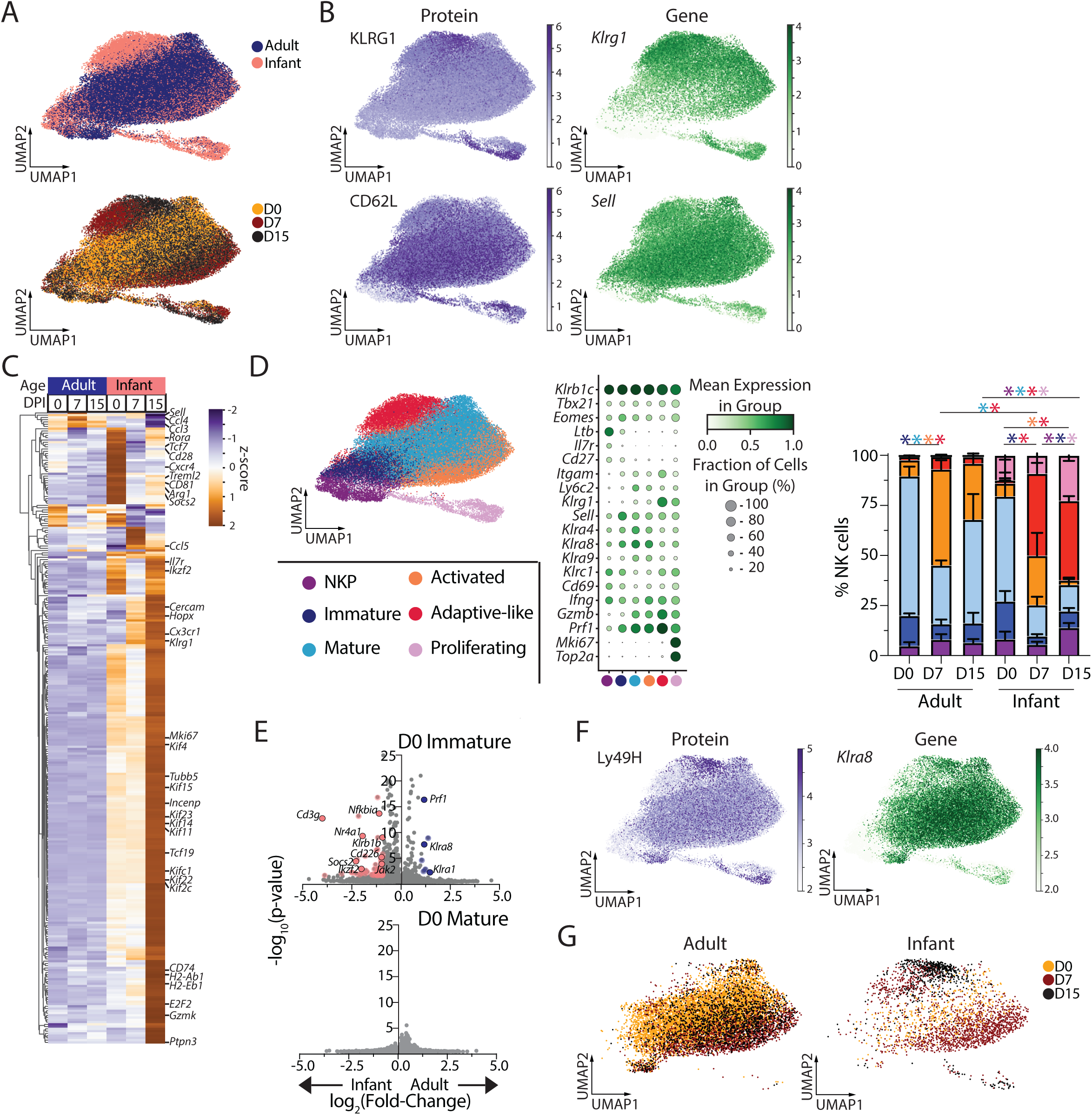
Distinct transcriptomic profile of infant and adult NK cells prior to and following MCMV infection. NK cells were isolated by magnetic enrichment from the spleens of naive and adult mice (D0) and 7 and 15 days post-MCMV infection (D7, D15) for CITE-seq profiling using the 10x genomics system. (A) UMAP embeddings of NK cells from top: adult (blue) and infant (pink) mice or bottom: by infection state D0 (gold), D7 (maroon), or D15 (black). (B) Expression of surface molecules bound by CITE-Seq antibodies (left) and corresponding gene expression (right) for KLRG1 (top) and CD62L (bottom). (C) Z-scored pseudobulk heatmap of the 100 most differentially expressed genes (p<0.05, log_2_(fold change)>1) for each group of enriched splenic NK cells from adult (blue) and infant (pink) mice at D0, D7, and D15 post-infection. (D) Leiden clustering (left) of adult and infant NK cells pre- and post-infection, with legend indicating cluster identity. Dot plot (middle) displaying the relative frequency and degree of expression for indicated genes that define subsets based on Leiden clusters. Proportion (right) of each NK cell subset for adult and infant mice pre- and post-infection. n = 4 recipients per age group per timepoint; *p<0.05 as assessed by two-way ANOVA with Bonferroni multiple comparisons test; asterisk color corresponds to the population that is significantly different. (E) Volcano plots displaying differentially expressed genes (pink/blue dots; p-value <0.05, log_2_(fold change)>1) between adult and infant prior to infection among immature (top) and mature (bottom) NK cells. (F) Expression of Ly49H bound by CITE-Seq antibodies (left) and corresponding gene expression (right) for *Klra8*. (G) UMAP embeddings of *Klra8*-expressing NK cells from adult (left) and infant (pink) mice by infection state D0 (gold), D7 (maroon), or D15 (black).

Following infection, infant NK cells were enriched in signatures associated with activation/ effector function (*Hopx*, *Gzmk*, *Socs2*, *Ptpn3*), cell migration (*Cxcr4*, *Cx3cr1*), antigen presentation (*CD74*, *H2-Ab1*, *H2-Eb1*, *CD81*), cell proliferation, and adaptive-like responses (*Sell*, *Klrg1*) compared to adult NK cells (**Fig. 3C**). These results reveal discrete gene expression profiles in infant compared to adult NK cells at baseline and following infection.

Leiden clustering of the NK cells (marked by expression of *Klrb1c*, *Tbx21*, and *Eomes*) revealed 6 discrete populations classified by canonical gene expression profiles (**Fig. 3D**). We identified NK cell precursors (NKP) expressing *Ltb*, *Il7r*, *Cd27* and lacking *Itgam* (CD11b_ while major NK cells subsets were delineated based on expression of *Cd27* and *Itgam* into immature (*Cd27*-expressing, *Itgam*-low-expressing) and mature (*Cd27*-non-expressing, *Itgam*-expressing) (**Fig. 3D**). Activated NK cells had increased expression of *Cd69* and *Ifng* relative to mature NK cells (*44*), while adaptive or memory-like NK cells expressed high levels of Klrg1/*Klrg1*and effector genes (*Ifng*, *Gzmb*, and *Prf1*) and low levels of CD62L (*20*, *23*). Additional clustering identified licensed NK cells, which interact with MHC Class I (*13*) (**fig. S3**).

The composition of NK cell subsets differed between infant and adult NK cells; infant NK cells had an increased frequency of proliferating cells and immature NK cells prior to infection (**Fig. 3D**). Prior to infection infant and adult NK cells exhibit similar transcriptome profiles, except for a few transcriptional differences between infant and adult immature NK cells (**Fig. 3E**). Following MCMV infection, however, adaptive-like NK cells predominated in infants while activated effector NK cells were the majority population in MCMV-infected adults for both total and Ly49H^+^ NK cells (**Fig. 3D,F,G; fig. S3**). Differentiation of adaptive NK cells exhibited distinct kinetics for infant Ly49H^+^ NK cells which acquired an activated profile starting at 7 DPI) followed by the establishment of an adaptive-like population at 15 DPI; while adult Ly49H^+^ NK cells also exhibit an activated profile at 7 DPI, they did not progress to adaptive-like cells (**Fig. 3G**). These CITE-seq data provide evidence for age-dependent trajectories of NK cell differentiation in responses to MCMV infection, and that the distinct responses are driven by age-specific cell fates.

### Distinct dynamics and responses of infant and adult MCMV-specific T cells

To address whether the distinct responses of infant and adult NK cells to MCMV infection occurred in the context of variable T cell responses, we investigated MCMV-reactive CD8^+^ T cells in two ways. First, we used peptide-MHC tetramers to identify MCMV-specific CD8^+^ T cell populations exhibiting conventional effector-to-memory transition (M45 epitopes) or followed inflationary kinetics through regular antigen encounter (M38 epitope) (*27*, *45*). The kinetics and magnitude (frequency and numbers) of virus-reactive M45 or M38-specific CD8^+^ T cell response to MCMV infection across sites was broadly similar in both infant and adult mice across all sites examined up to 60 DPI (**Fig. 4A,B**; **fig. S4**). Because the infant mice were maturing to adults over the time course, the involvement of new adult-like T cells could affect the results. Therefore, we implemented a co-adoptive transfer model to allow individual tracking of infant and adult-derived T cell responses in the same mouse.

**Figure 4.**
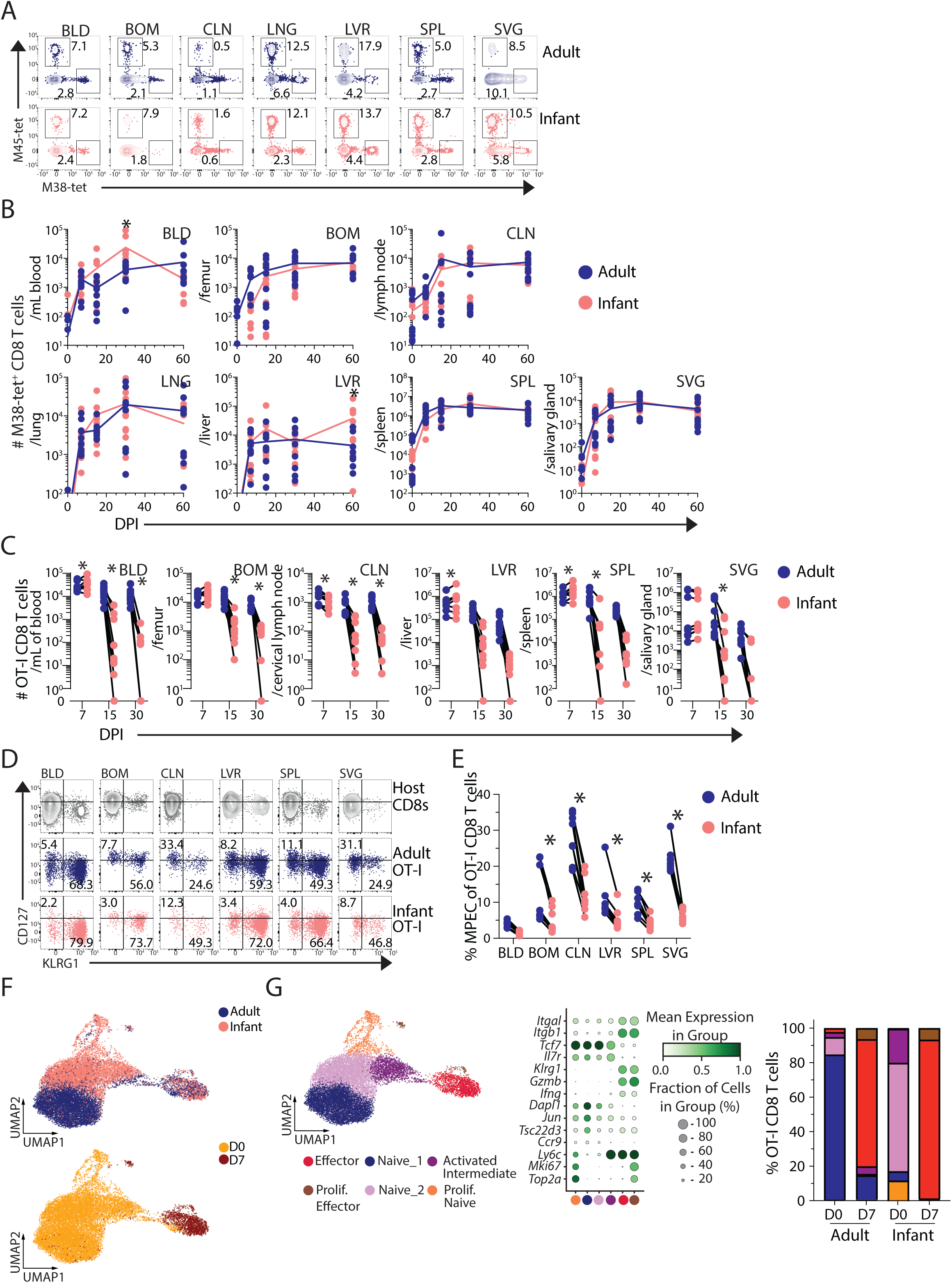
Infant T cells have intrinsically reduced memory generation to MCMV infection. (A) Representative gating of MCMV-tetramer^+^ M45 or M38 CD8^+^ T cells. Gates are based on uninfected (DPI 0) mice. (B) Number of M38-specific ‘inflationary’ CD8^+^ T cell populations across tissues from adult (blue) and infant (pink) mice. (C) Experimental design in fig. S5A. Number of adult (blue) and infant (pink) OT-I CD8^+^ T cells from MCMV-OVA infected hosts at indicated DPI. (D) Representative flow cytometry plots showing frequency of MPEC (CD127^hi^ KLRG1^-^) or SLEC (CD127^lo^ KLRG1^+^) of total recipient CD8^+^ T cells (grey), or adult (blue) or infant (pink) OT-I CD8^+^ T cells at 7 DPI. (E) Frequency of MPEC adult (blue) and infant (pink) OT-I CD8^+^ T cells from respective tissue. n = 10 mice per time-point per group. Lines connecting data points indicate paired transfers. * p-value < 0.05 assessed by two-way ANOVA with Bonferroni multiple comparisons test. Data are combined from N=2 independent experimental replicates. Tissue Abbreviations: blood (BLD), bone marrow (BOM), cervical lymph node (CLN), lung (LNG), liver (LVR), spleen (SPL), and salivary gland (SVG). OT-I CD8^+^ T cells were isolated from the spleen of uninfected infant and adult OT-I donor mice (D0) and from C57Bl/6 recipient spleens 7 days post-transfer/ MCMV infection. (F) UMAP embedding of OT-I CD8^+^ T cells from top: adult (blue) and infant (pink) donors or bottom: by infection status D0 (gold) and D7 (maroon) post-infection. (G) Leiden clustering (left) of adult and infant OT-I CD8^+^ T cells pre- and post-infection, with legend indicating cluster identity. Dot plot (middle) displaying the relative frequency and degree of expression for indicated genes that define subsets based on Leiden clusters. Proportion (right) of each OT-I CD8^+^ T cell subset for adult and infant mice pre- and post-infection.

In the co-transfer model, equal numbers of T cell receptor-transgenic (TCR-tg) OVA-specific T cells from congenically distinct infant (Thy1.1^+/+^ CD45.2^+/+^) and adult (Thy1.2^+/+^ CD45.2^+/+^) OT-I mice (**fig. S5A,B**) were co-transferred into adult (CD45.1^+/+^) hosts, which were infected a day later with recombinant MCMV expressing ovalbumin (MCMV-OVA) (*28*) and followed from 7-30 DPI (**fig. S5A**). At 7 DPI, the number of infant-derived OT-I CD8^+^ T cells exceeded adult-derived cells in the blood, spleen, and liver indicating a more robust acute response; by contrast, at later times (15, 30 DPI), the number of infant relative to adult OT-I CD8^+^ T cells was significantly lower in all sites with negligible infant OT-I CD8^+^ T cells present at 30 DPI (**Fig. 4C**). These results show that infant T cells undergo a robust expansion in multiple sites of infection but subsequently contract and fail to persist as memory T cells. This rapid loss of MCMV-specific infant OT-I cells during acute infection correlated with higher frequencies of short-lived effector cells (SLECs; CD127^lo^ KLRG1^+^) and reduced memory precursor effector cells (MPECs; CD127^hi^ KLRG1^-^) (*46*, *47*) compared to adult OT-I cells at 7 DPI (**Fig. 4D, E**). These findings show altered T cell fates between infants and adults manifested at relatively early times post-infection.

CITE-seq profiling of infant and adult OT-I CD8^+^ T cells prior to transfer and 7 DPI in the co-transfer model, revealed major transcriptional differences based on origin (Infant/Adult) and activation state (D0 and D7 post-infection) (**Fig. 4F, fig. S5B-C**, and **Table S2**). At baseline, there were three naïve-like OT-I subpopulations for infant compared to one for adult T cells **(Fig. 4G**). By 7 DPI, the majority of infant and adult-derived OT-I cells exhibited overlapping effector profiles (*Ifng, Gzmb*, *Itgal, Itgb1*) resembling SLECs (**Fig. 4G**), while adult OT-I also contained a population that transcriptionally and phenotypically resembled MPECs (*46*, *48*) (**fig. S5D, E**). This infant SLEC bias at 7 DPI was also found for M45- and M38- tetramer^+^ cells derived from direct infection of infant compared to adult mice (**fig. S6**). Together, these results demonstrate increased memory formation by adult compared to infant T cells, which are biased to effector differentiation and ultimately cell death.

### An age-dependent switch in memory fates by NK and T cells

The differences between infant versus adult T and NK cells to form memory could arise from intrinsic features of neonatal lymphocytes that shift when mice became adults or progressive changes in cell fate with age. To follow the differentiation fate of OT-I CD8^+^ T cells we set up timed pregnancies to yield OT-I pups on a weekly basis with conserved paternity.

Splenic OT-I CD8^+^ T cells from mice aged 10- 45 DPN were co-transferred with congenic OT-I CD8^+^ T cells from adult mice into adult recipients, which were subsequently infected with MCMV-OVA and memory generation was assessed at 30 DPI (**Fig. 5A**). We found similar yields of memory OT-I CD8^+^ T cells in all recipient mice indicating a similar infection response by all groups (**Fig. 5B**); however, the young:adult ratio of OT-I CD8^+^ T cells differed by age group. The frequency of infant or weanling OT-I CD8^+^ T cells increased relative to adult OT-I from 10-45 days post-natal age (**Fig. 5C, D**). This result provides evidence for progressive differentiation of infant T cells to become adult-like in their capacity to form memory and establish protective immunity, consistent with findings from human T cells (*49*).

**Figure 5.**
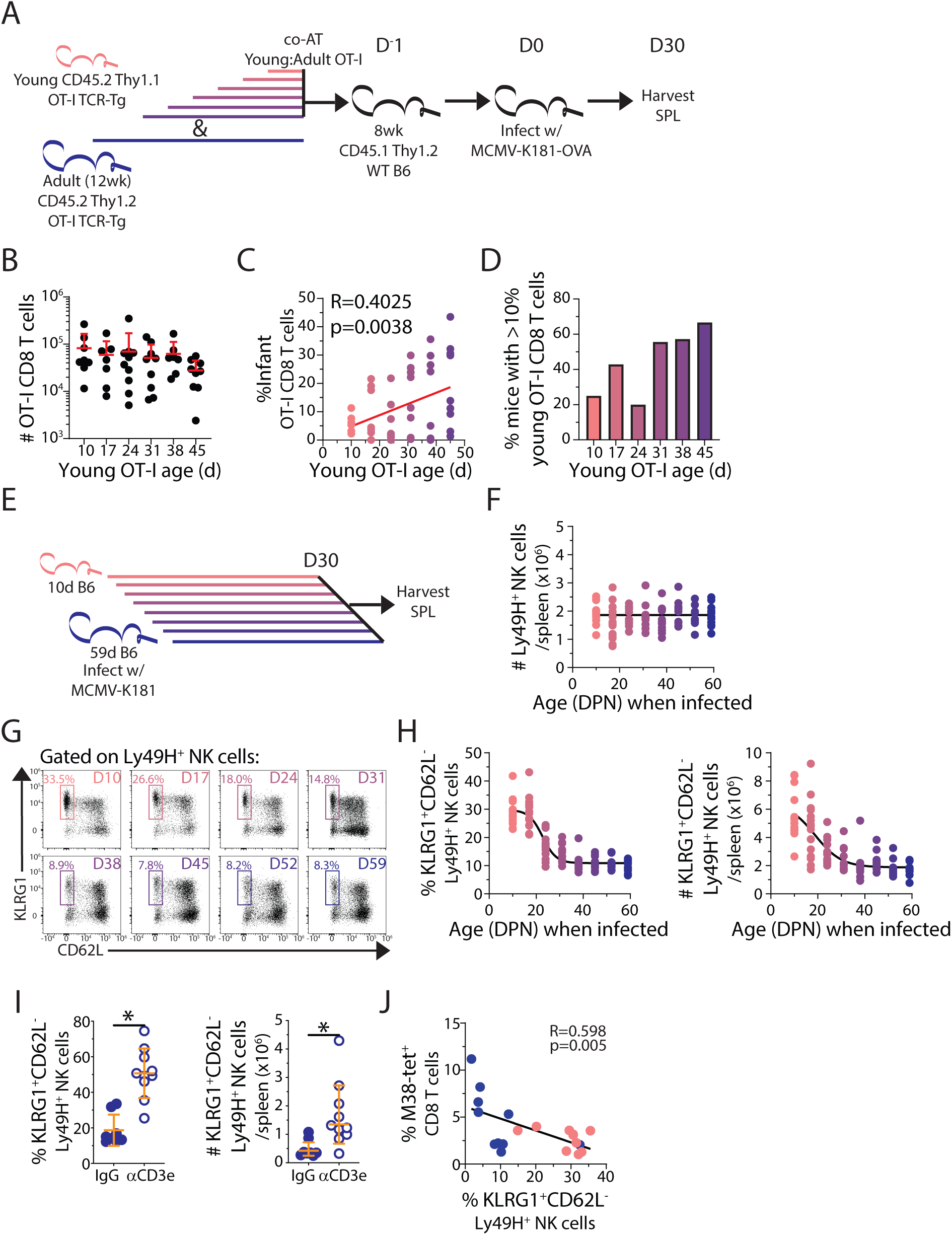
Age-dependent changes in memory potential for T and NK cells. (A) Experimental design for panels B-E. (B) Total number of recovered OT-I CD8^+^ T cells and (C) frequency of recovered young OT-I CD8^+^ T cells in spleen at D30 post-infection. n = 7-10 per group. Red line: linear regression with significance and fit determined using Spearman correlation. (D) Percentage of individual host mice with >10% recovered young OT-I CD8^+^ T cells in spleen at DPI 30. (E) Experimental design for panels G-H. (F) Total number of Ly49H^+^ NK cells per spleen at D30 post-infection. (G) Representative gating of adaptive-like (KLRG1^+^ CD62L^-^) NK cells. (H) Frequency (left) and number (right) of splenic adaptive-like (KLRG1^+^ CD62L^-^) NK cells of total Ly49H^+^ NK cells at 30 DPI of mice infected at indicated ages. n = 10 individual mice per group. (I) Frequency (left) and number (right) of adaptive-like (KLRG1^+^ CD62L^-^) Ly49H^+^ NK cells in spleens at 15 DPI of mice treated with control IgG or αCD3-depleting antibody. n = 8-10 individual mice per group. * p-value < 0.05 as assessed by t-statistic. (J) Linear regression with Spearman correlation of the frequency of adaptive-like (KLRG1^+^ CD62L^-^) Ly49H^+^ NK cells and the frequency of the M38-tetramer^+^ ‘inflationary’ CD8^+^T cells at 60 DPI. All data are combined from N= 2 independent experimental replicates.

To determine whether the age-dependent difference in MCMV-reactive NK cell response also occurred during a similar maturation window as T cells, we infected littermate cohorts with MCMV on a weekly basis between 10-60 DPN and assessed the adaptive NK cell response 30 DPI (**Fig. 5E**). Importantly, by 30 DPI the number of total Ly49H^+^ NK cells was equivalent in all experimental groups (**Fig. 5F**). However, the composition of Ly49H^+^ NK cells over age resulted in progressively lower frequencies and numbers of adaptive-like (KLRG1^+^ CD62L^-^) NK cells (**Fig. 5G, H**). The reduced frequencies of adaptive-like NK cells with age was coincident with the age-associated increase in memory capacity by CD8^+^ T cells, revealing a coordinate age-dependent shift in differentiation fate from NK cell- to T cell-memory.

To establish whether T cell competition limits the development of adaptive-like NK cells, we performed antibody-mediated T cell depletion in adult mice relative to IgG-treated controls at the peak (15 DPI) of adaptive-like NK cell expansion (**Fig. 5I**). T cell-depleted mice had a more robust expansion by proportion and numbers of adaptive-like NK cells (**Fig. 5I**) indicating that competition with T cells limits adaptive-like NK cell responses. Further, within intact mice, an inverse relationship was present among the M38-tetramer^+^ ‘inflationary’ population of CD8^+^ T cells and the formation of adaptive-like NK cells (**Fig. 5J**), suggesting a competitive dynamic between the memory fates of NK cells and CD8^+^ T cells that may contribute to the enhanced adaptive-like NK cell response in infants.

### Age influences adaptive-like human NK cell phenotypes and abundance to HCMV

Our findings of enhanced differentiation of adaptive-like NK cells in early life prompted examination of whether human NK cells exhibit analogous age-dependent fates in HCMV infection. Human NK cells in adults comprise a CD56^dim^ “mature” or effector-like subset found predominantly in blood, bone marrow (BOM), spleen, and lungs along with a CD56^bright^ “immature” subset that are localized to lymph nodes (LNs) and gut and can be tissue resident (*50*). HCMV-reactive NK cells in human blood are distinguished by NKG2C and CD57 expression (*17*, *51*, *52*), though signatures for adaptive-like NK cells across human tissues have not been defined. Here, we used our extensive human tissue resource whereby we obtain blood and multiple lymphoid and mucosal tissue from human organ donors of all ages to investigate HCMV-associated NK responses between children and adults and over age.

To identify putative adaptive-like NK cells associated with HCMV infection in human blood and tissues, we interrogated our recent CITE-seq database of immune cells across 10 sites of 24 donors (8- HCMV^-^, 16- HCMV^+^)(*53*). We annotated >130,000 Natural Killer and Innate Lymphoid Cells (NK/ILCs) and used single cell Hierarchical Poisson Factorization (scHPF) analysis (*54*, *55*) to identify co-expressed gene module(s) enriched in HCMV^+^ individuals (**Fig. 6A**). A single gene module associated with HCMV infection was demarcated by genes that canonically define the HCMV-specific NK cell responses (*KLRC2* [NKG2C] and *B3GAT1*[enzyme that produces CD57]) (*17*) and also included genes encoding activating/inhibitory receptors (*LAG3*, *KLRC3*, *KLRG1*, *MCOLN2*), effector molecules/cytokines (*GZMH*, *CCL5*, *PROK2*), and HLA class II (*HLA-DRB5*, *HLA-DRB1*, *HLA-DRA*) (**Fig. 6A, Table S3**).

**Figure 6.**
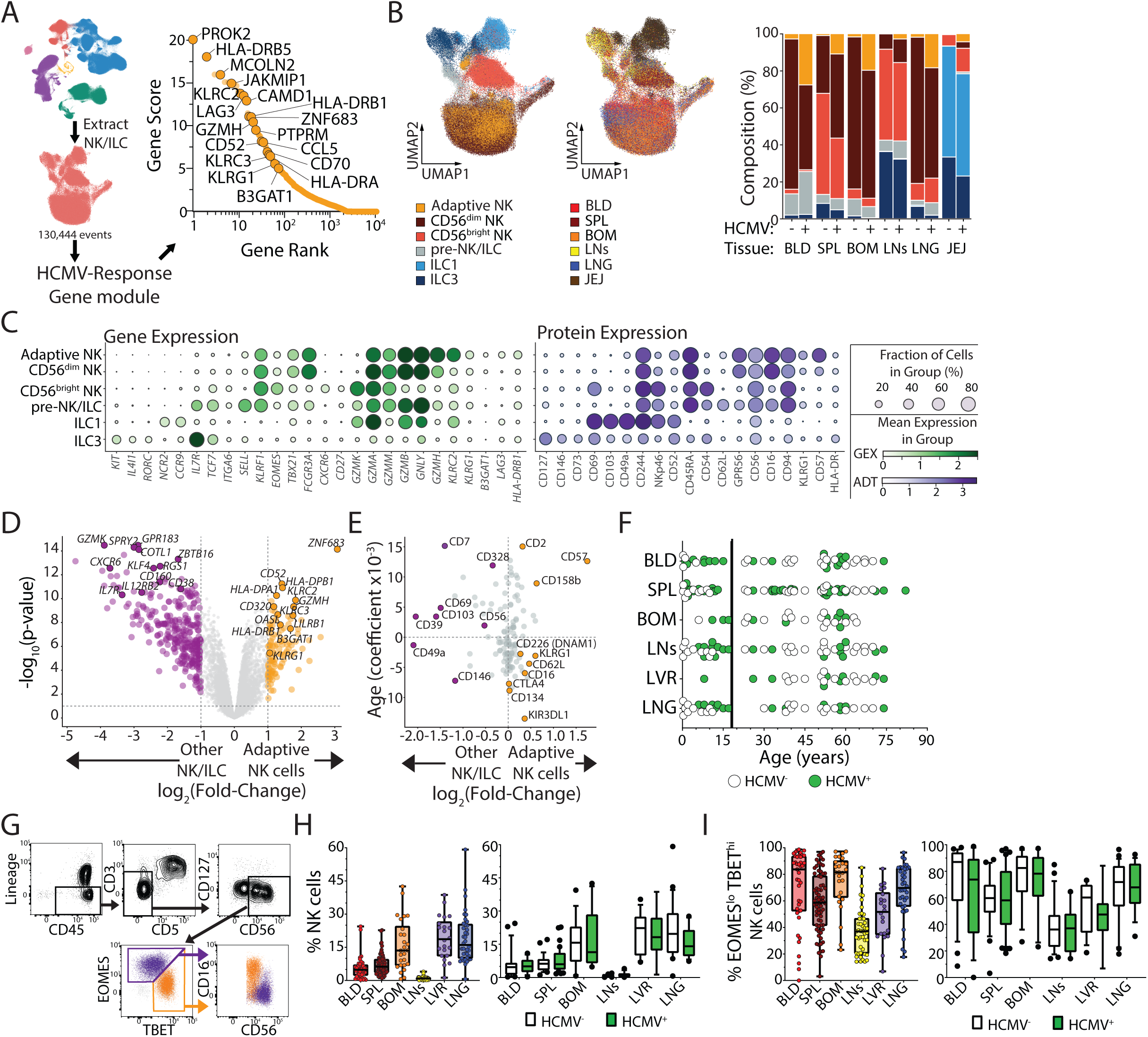
HCMV infection elicits a discrete gene expression module by NK cells. (A) HCMV-specific adaptive-like NK cell gene module derived from Ref. 55. (B) UMAP embedding of NK/ILC cell populations (left; Adaptive-like- gold, CD56^dim^- maroon, CD56^bright^-red, pre-NK/ILC- grey, non-NK ILC1- light blue, ILC3s- dark blue) and tissue distribution (middle; blood- red, spleen- maroon, bone marrow- orange, lymph nodes-yellow, lung- blue, and jejunum- brown). Quantification of NK/ILC populations (right) in indicated tissues from HCMV^-^and HCMV^+^ individuals. (C) Dot plots displaying the frequency and relative expression of indicated genes (left; green) and proteins (right; purple) corresponding to NK/ILC subset identities. (D) Volcano plot displaying differentially expressed genes (purple/gold dots; adjusted p-value <0.05, log_2_(fold change)>1) between adaptive-like NK cells and all other NK/ILC subsets. (E) Quadrant plot for adaptive-like NK cell protein expression relative to all other NK/ILC subsets and the influence of age (age coefficient) on that protein expression. (F) Donor (N= 76) tissue distribution by age from pediatric [<18 years old; N= 31] and adult [≥18 years old; N= 45] (blue) delineated by HCMV-serostatus (HCMV^-^-white, HCMV^+^- green). (G) Representative gating of NK cells and the association of EOMES/TBET expression with CD56^bright^ and CD56^dim^ NK cells. (H) Frequency of NK cells among CD45^+^ lymphocytes from all donors (left) and stratified by donor HCMV-serostatus (right). (I) Identical analysis as panel I for EOMES^lo^ TBET^hi^ [i.e., CD56^dim^] expression among NK cells. Data are combined from 10 batched experiments assessing 76 unique donors. Tissue Abbreviations: blood (BLD), spleen (SPL), bone marrow (BOM), lymph nodes (LNs; hepatic and lung), liver (LVR), lung (LNG), and jejunum (JEJ)

This adaptive-like signature was used to annotate our NK/ILC CITE-seq dataset along with known signatures for CD56^bright^, CD56^dim^, and different ILC populations using the MMoChi algorithm (*56*). Adaptive-like NK cells were a subset of CD56^dim^ cells distinct in profile and tissue distribution from CD56^bright^ and ILC cells; adaptive-like cells were abundant in the blood, BOM, spleen and lung, while CD56^bright^ and ILC were found mostly in LN, spleen, and intestine (**Fig. 6B**). Importantly, the proportion of adaptive-like NK cells was higher in the blood, spleen, BOM, and lungs of HCMV^+^ compared to HCMV^-^ donors (**Fig. 6B**). Gene and protein expression of the classified subsets showed that adaptive-like NK cells differed from other CD56^dim^ NK cells in upregulated expression of *KLRC2*, *GZMH*, *LAG3* genes, CD52 protein expression, and *KLRG1*/KLRG1, *B3GAT1*/CD57 gene/protein expression (**Fig. 6C**). Relative to all other NK/ILCs, adaptive NK cells exhibited decreased expression of genes associated with progenitors (*IL7R*, *KLF4*, *ZBTB16*) (*57*) and lower surface expression of molecules associated with tissue residency (CD49a, CD39, CD103, CD69) (*50*, *58*, *59*) (**Fig. 6D**). Across age (20-75yrs), adaptive NK cells showed increased expression of CD57, a marker of replicative senescence, while other markers (KLRG1, CD62L, CD226) did not change (**Fig. 6E**). Together, these findings reveal an adaptive-like NK cell subset that is enriched in HCMV^+^ individuals that changes across a broad age range in adults.

Based on the CITE-seq profiles, we designed a high dimensional flow cytometry panel to comprehensively profile HCMV-associated NK cell responses across blood, tissues, and age. The cohort of 76 organ donors spanned 3 months to 82 years of age, of whom 59% were HCMV^+^ and comprised 31 children (<18yrs) and 45 adults with similar representation for blood and most tissues (Spleen, LNs lungs) except for BOM and liver where pediatric samples were lacking (**Fig. 6F, fig. S7**, and **Table 4**). NK cells (CD45^+^CD14/CD19/CD3-CD127^-^CD56^+^, see **fig. S1B** for gating strategy) were further subdivided by expression of transcription factors EOMES and TBET into EOMES^hi^ TBET^lo^ and EOMES^lo^ TBET^hi^ cells (**Fig. 6G**) representing CD56^bright^ CD16^-^ (immature) and CD56^dim^ CD16^+^ (mature) classifications, respectively(*60*, *61*). The proportion of total and EOMES^lo^ TBET^hi^ (mature) NK cells varied by tissue but not HCMV status or age. Notably, BOM liver, and lungs had higher frequencies of total NK cells compared to blood and spleen while LNs had very low frequencies overall (**Fig. 6H**; **fig. S8A**). By contrast, mature NK cells were enriched in all sites except the LNs (**Fig. 6I**; **fig. S8A**).

We defined HCMV-reactive NK cells based on expression of NKG2C along with its co-receptor CD94(*62*). NKG2C^+^ CD94^+^ NK cells were significantly more abundant in the spleen, liver, and lungs (and an increased trend in the BOM) of HCMV^+^ compared to HCMV^-^ donors (**Fig. 7A**) and were equally abundant in HCMV^+^ pediatric and adult tissues. We assessed the expression of both CD57 and KLRG1 by HCMV-reactive NKG2C^+^ CD94^+^ cells to identify adaptive-like subsets (**Fig. 6F**). CD57 was specifically expressed by NKG2C^+^ NK cells in HCMV^+^ adult donors, pediatric HCMV-reactive NK cells did not express CD57 (**Fig. 7B**).

**Figure 7.**
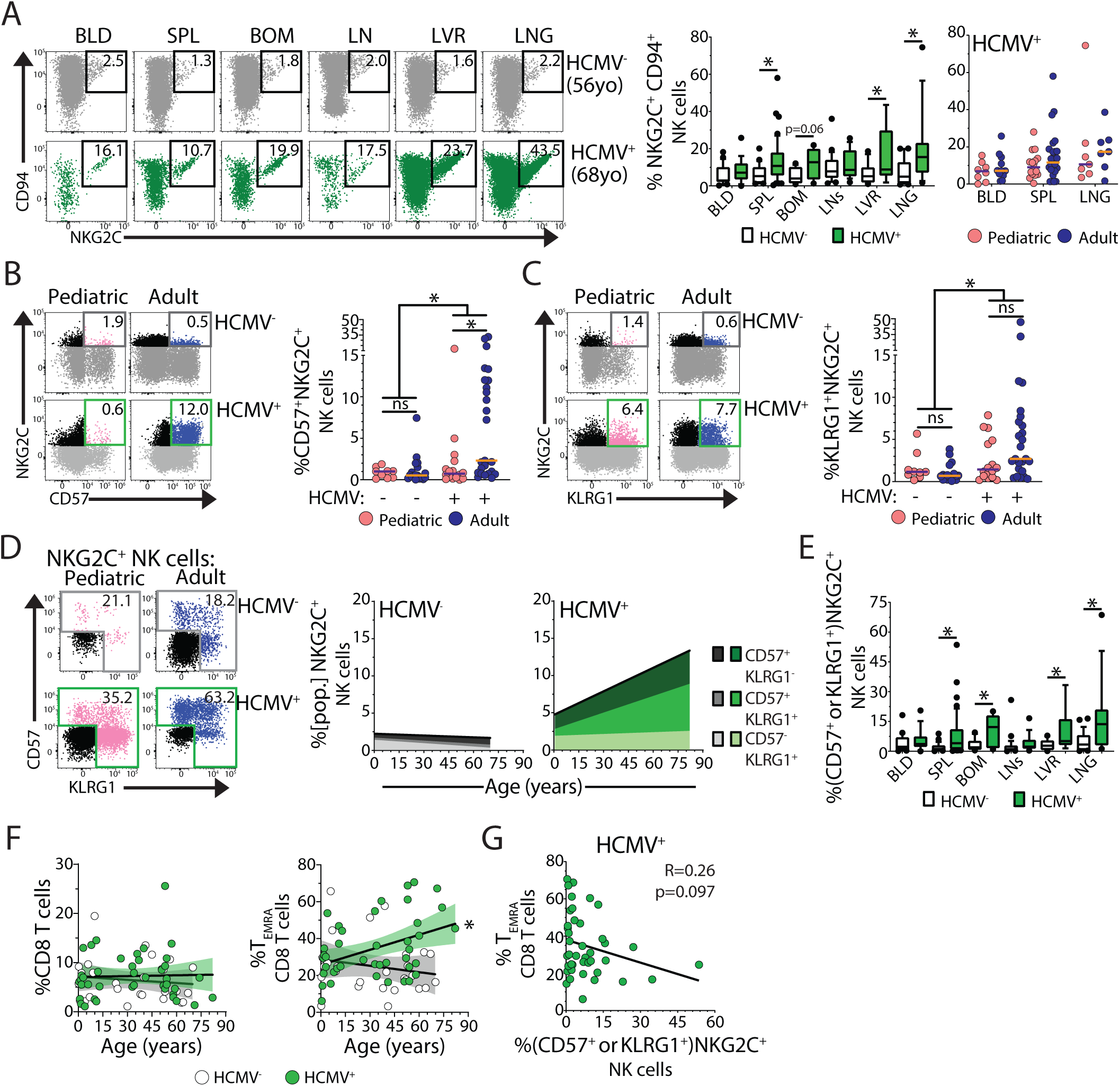
Distinct frequencies and magnitude of HCMV-reactive adaptive-like NK cells over age. (A) Representative gating (left), frequency of NKG2C-expression (NKG2C^+^ CD94^+^) among NK cells stratified by donor HCMV-serostatus (middle; HCMV^-^- white, HCMV^+^- green) and stratified by age cohort among HCMV^+^ individuals (right; pediatric- pink, adult- blue) in indicated tissues. * p-value < 0.05 assessed by two-way ANOVA with Bonferroni multiple comparisons test. (B) Representative gating (left), frequency of splenic CD57^+^ NKG2C^+^ NK cells (right) from HCMV^-^ and HCMV^+^ pediatric (pink) and adult (blue) donors. * p-value < 0.05 assessed by two-way ANOVA with Bonferroni multiple comparisons test. (C) Identical analysis as panel C for KLRG1^+^ NKG2C^+^ NK cells. (D) Representative gating and stacked linear regressions for subsets of splenic CD57- and KLRG1-expressing NKG2C^+^ cells among NK cells over age in HCMV^-^ (grey) and HCMV^+^ (green) donors. (E) Frequency of adaptive-like ([CD57 and/or KLRG1]-expressing NKG2C^+^) cells among NK cells in indicated tissues from HCMV^-^(white) and HCMV^+^ (green). (F) Linear regression with Spearman correlation of splenic CD8^+^ T cells (left) and CD8^+^ T_EMRA_ cells (right) stratified by HCMV-serostatus (left; HCMV^-^- white, HCMV^+^- green). (G) Linear regression with Spearman correlation of splenic CD8^+^ T_EMRA_ cells versus adaptive-like NK cells in HCMV^+^ donors. Data are combined from 10 batched experiments assessing 76 unique donors. Tissue Abbreviations: blood (BLD), spleen (SPL), bone marrow (BOM), lymph nodes (LNs; hepatic and lung), liver (LVR), and lung (LNG).

However, KLRG1^+^ NKG2C^+^ NK cells were enriched in HCMV^+^ compared to HCMV^-^ donors for both pediatric and adult donors (**Fig. 7C**), indicating that this marker can delineate early adaptive-like NK cells as in mice. Over age, there was an accumulation of CD57^+^ KLRG1^+^ and CD57^+^ KLRG1^-^ HCMV-reactive NK cells while CD57^-^ KLRG1^+^ subsets persisted in stable frequencies (**Fig. 7D**; **fig. S8B, C**). Interestingly, CD57^-^ KLRG1^+^ cells were more abundant in lymphoid tissue while CD57^+^KLRG1^+/-^ cells prevailed in the major tissue reservoirs like liver and lung (**fig. S8D**). Moreover, CD57 and/or KLRG1 expression by NKG2C^+^NK cells was enriched in the spleen, BOM, liver and lungs but not in blood of HCMV^+^ individuals (**Fig. 7E, fig. S8E**), indicating that the majority of HCMV-reactive adaptive-like NK cells are maintained in tissues rather than circulation.

We further compared the dynamics of adaptive-like NK cells with HCMV-driven T cell responses to assess whether they were interrelated as we found in MCMV infection. As in mice, CD8^+^ T cells in humans exhibit high frequency responses to HCMV and HCMV-reactive T cells are predominantly maintained as terminally differentiated effector or TEMRA cells that circulate through blood, BOM, and spleen (*30*, *31*). Similar to prior reports (*31*, *53*, *63*), while the frequency of total CD8^+^ T cells was stable over age regardless of HCMV-serostatus, CD8^+^ T_EMRA_ cells increased substantially over age in the spleen and blood of HCMV^+^ but not HCMV^-^individuals (**Fig. 7F** and **fig. S8F-H**). We further found an inverse association in the proportion of splenic HCMV-reactive adaptive-like NK cells and CD8^+^ T_EMRA_ cells (**Fig. 7G**). Together, these results indicate that the human adaptive-like NK cell response to HCMV is informed by age, tissue, and competition with CD8 T cells, qualities shared with responses to MCMV.

## DISCUSSION

CMV infection is prevalent globally, with many infections occurring during infancy and childhood, though it is not known whether the age of initial infection affects long-term immunity. Here, we investigated the immune response to CMV infection from infancy through adulthood using a mouse MCMV infection model and human samples, revealing age-dependent control of NK cell differentiation, which compensates for limited T cell memory establishment in early life. Adaptive-like NK cells were preferentially formed during infancy when they seeded tissue reservoirs, and their proportion declined with age coinciding with an increase in MCMV-reactive T cell memory. In humans, early adaptive-like NK cells were found in HCMV seropositive children, where they exhibited biased distribution in tissues, underwent age-dependent phenotypic changes, and competed with T cells for niche occupancy. Together, our findings reveal that NK cell memory to CMV infection is established in early life and maintained in tissue reservoirs providing a distinct type of long-term tissue protection.

It is well established that in humans, NK cells play a key role in control of persistent infections such as HCMV, based on studies of individuals lacking NK cells (*8*, *10*, *64*). NK cells can establish adaptive-like responses to infections with CMV being the most well-studied, given the identification of CMV-reactive populations (Ly49H^+^ in mice and NKG2C^+^ in humans) that exhibit enhanced effector responses to infectious re-challenge (*17*, *20*, *65*). Here, we show that CMV infection during infancy or childhood results in predominant expansion of NK cells that exhibit transcriptional and phenotypic features of adaptive-like NK cells and are maintained in multiple tissues for both MCMV-infected mice and HCMV-infected humans. In mice, the magnitude of the adaptive-like NK cell response was independent of viral load and could be enhanced by depleting T cells, indicating that age and the host immune environment influence NK memory formation. Previous studies found that blood NK cells from newborns with congenital HCMV infection were more activated and effector-like due to in utero activation (*66*, *67*) and could also become hyporesponsive (*68*). Our findings show that postnatally, infant NK cell responses to CMV are distinct from both adult and perinatal/fetal infection, with infancy/childhood being a prime window for the formation of adaptive-like NK cells to shape the NK repertoire for lifelong immunity.

In contrast to the enhanced memory formation of infant NK cells, the infant CD8^+^ T cell response to MCMV was marked by extensive effector cell differentiation at the expense of memory cells. The overall reduced memory fate of infant T cells is consistent with results in influenza and Listeria infection models by our group and others (*32–34*), due in part to the enhanced signaling sensitivity by infant T cells (*32*). Thus, the elevated development of adaptive-like NK cells during infancy compensates for the limited T cell memory capacity, and this progressively shifts with age as T cells acquire memory capacity. This shift in memory propensity by NK and T cells occurs during immune reconstitution following hematopoietic stem cell transplantation where NK cell expansion precedes CD8^+^ T cell responses (*69*). The inverse association between CD8^+^ T_EMRA_ cells and adaptive-like NK cells in the spleen indicates that niche occupancy may function during CMV persistence. We propose that individuals having substantial frequencies of HCMV-reactive NK cell memory likely experienced early life infection. In this way the relative proportions of innate versus adaptive memory to an infection may represent a time stamp of the initial infection environment.

We identified a unifying signature for HCMV-associated adaptive-like NK cells across tissues through CITE-seq profiling. This adaptive-like subset, found primarily among infant NK cells, shares transcriptional features of human adaptive NK cells recently reported (*70*). Indeed, we observe that the abundance of HCMV-reactive adaptive-like NK cells increases with host age and this increased population expresses CD57, indicating repeated stimulation or replicative senescence. In HCMV infection CD8^+^ T_EMRA_ cells also exhibit increased frequencies with age and acquire CD57 expression (*31*). Interestingly, adult adaptive-like NK cells transcriptionally resemble CD8^+^ T_EMRA_ cells (*71*). In this light, predominant CD57^+^ KLRG1^+/-^ cells that designate adaptive-like NK cells found in the blood (*17*) may derive from KLRG1^+^CD57^-^ NK cells stably maintained in tissues. Identifying the relationships between the different adaptive-like NK cell populations may facilitate targeting these populations for long-term immune memory.

In conclusion, our findings provide rationale for early life vaccination against CMV and other infections to generate durable NK cell-mediated immune memory, which can be more versatile and elicit broader protection than T cells (*72*, *73*). The ability of NK memory to populate systemic tissue reservoirs is important to contain and prevent infection dissemination during CMV persistence as well as potential reactivation that can occur due to transient or prolonged immunosuppression. Understanding the niche competition factors globally and locally will also be important for tuning the responses of the targeted memory population for NK versus T cell memory.

## MATERIALS AND METHODS

### KEY RESOURCES TABLE

A list of antibodies, reagents, and software used to effectuate the following methodologies can be found in **Table S5**.

## RESOURCE AVAILABILITY

### Lead contact

Further information and requests for reagents should be directed to the lead author.

### Materials availability

This study did not generate new unique reagents.

## EXPERIMENTAL MODELS AND SUBJECT DETAILS

### Animals and ethics statement

Mice were housed and bred in specific pathogen-free conditions in the animal facilities at Columbia University Irving Medical Center (CUIMC). Male and female CD45.1 [B6.SJL-Ptprca Pepcb/BoyJ], Thy1.1 [B6.PL-Thy1^a^/CyJ], and OT-I [C57BL/6-Tg(TcraTcrb)1100Mjb/J] were purchased from Jackson Laboratory. Thy1.1 and OT-I mice were bred to generate Thy1.1 congenic hosts. All infant mice were used at 9-10 days of age, and adult mice were used at >8 weeks old. Infections were performed in biosafety level 2 biocontainment animal facilities. All studies utilized both male and female mice. All animal studies were approved by and performed in compliance with Columbia University Institutional Animal Care and Use Committee.

### Human samples

Blood and tissue samples were obtained from brain-dead human organ donors through collaborations with organ procurement organizations throughout the US. Samples from adult organ donors were obtained through a long-standing research protocol with LiveOnNY, the organ procurement organization for the New York metropolitan area as described (*74*, *75*).

Samples from infant and pediatric donors were obtained as part of the Human neonatal development-immunology (HANDEL-I) program based at the University of Florida as described (*49*, *76*). The use of organ donors does not qualify as human subjects research because samples are obtained from deceased and not living individuals, as confirmed by the Columbia University Institutional Review Board.

### Viruses, virus titrations and infections

Unless otherwise stated, mice were infected with 10^4^ plaque forming units (PFU) of MCMV-K181 or recombinant MCMV-OVA (*29*) administered intraperitoneally (i.p.). Viral stocks were produced by infecting M2-10B4 cells at a multiplicity of infection (MOI) 0.01. Upon visible cytopathic effect, virus was purified from supernatant and cell lysate by ultracentrifugation. Viral titrations were determined by standard plaque assay using M2-10B4 cells as previously described (*77*).

### *In vivo* NK and T cell depletions and morbidity evaluation

NK and T cell depletion was achieved by intraperitoneal (*i.p.*) injection with 300 μg of anti-mouse NK1.1 depleting antibody (PK136) (Bio-X-Cell) or 300μg anti-CD3ε depleting antibody (145-2C11) (Bio-X-Cell) at 3 days prior to infection, on the day of infection, and 7 days post infection, respectively. Isotype controls were performed at the same time-points with *i.p.* injection of 300μg of mouse IgG2a (C1.18.4) or 300μg Armenian Hamster IgG (PIP) (Bio-X-Cell), respectively. Mice were weighed every other day to evaluate for morbidity.

### Determination of MCMV viral load by quantitative polymerase chain reaction (qPCR)

To evaluate viral burden in acute infection, infant or adult tissues from infected mice were harvested at indicated timepoints. Tissues were homogenized using Qiagen Tissuelyser II (Qiagen) and DNA was extracted using Qiagen DNeasy Blood & Tissue Kit (Qiagen) per manufacturer protocols. qPCR for MCMV viral loads were performed using SsoAdvanced Universal SYBR Green Supermix® (Bio-Rad) on CFX384 Opus system (Bio-Rad). Cycling conditions were as follows: polymerase activation 2 min at 98 °C, followed by 40 cycles of denaturation for 15s at 98 °C, annealing/extension for 30s at 60 °C. Specificity of amplified sequences was validated by inspection of melting curves. Each individual sample for qPCR was ran in triplicate. Viral burden was quantified as fold change of cycle threshold (C_t_) values: 2^(C_t_ *M45*- C_t_ *Actb*) Primer sequences: M45 5’-TGTCTGTCGACGCACATC-3’and 5’-CCTTCTACGACGACTCGG-3’; *Actb* 5’-GCCAACTTTACGCCTAGCGT-3’ and 5’-ACCGGCTCATCAAATGCC-3’.

### Murine tissue processing for flow cytometry

Tissues were harvested at indicated days post-infection and processed to generate single-cell suspensions. Briefly, liver, lung, and salivary gland were dissociated using gentleMACS lung protocol (Miltenyi) and digested for 1 hour in digest media (RPMI 1640 medium containing collagenase D (Sigma-Aldrich), deoxyribonuclease (Sigma-Aldrich), trypsin inhibitor (Sigma-Aldrich)), in a 37 °C shaking incubator. Organ digests were disrupted through a 100 µm strainer, suspended in 25% solution of Percoll (Sigma) in complete media (IMDM (1X) (Gibco) supplemented with 10% FBS (GeminiBio) and 1% PSQ (100 U/ml penicillin, 100 μg/ml streptomycin, 2 mM L-glutamine, GeminiBio)), then centrifuged and supernatant aspirated. Lymph nodes and spleen were disrupted through a 100-um strainer (Corning) in complete media. Femurs were flushed with 30G syringe containing ice-cold complete media to extract bone marrow. Red blood cells were lysed in all tissue sites using ammonium-chloride-potassium (ACK) lysis buffer (Gibco) for 5 min before addition of complete media to neutralize the reaction. Cells were then washed with complete media and counted using Solution 13 (AO+DAPI) on NucleoCounter-3000 (Chemometec).

### Cell analysis by flow cytometry

A list of all antibodies used for flow cytometry are shown (**Table S3**). Following tissue processing, single cell suspensions were plated in FACs buffer (PBS (Corning) + 2mM EDTA (Corning) + 5% FBS (GeminiBio)). Cells were stained with surface markers for 30 min at 4 °C. To evaluate transcription factor expression and intracellular markers, cells were then fixed and permeabilized with Foxp3 fix/perm solution (Tonbo) before staining for 30 min at 4 °C. Dead cells were excluded using Zombie NIR Fixable Viability Kit (Biolegend) according to manufacturer’s instructions. For detection of MCMV-specific CD8^+^ and CD4^+^ T cells, MHC class I tetramers H-2D(b) MCMV M45 985-993 HGIRNASFI and H-2K(b) MCMV M38 316-323 SSPPMFRV were obtained from National Institutes of Health (NIH) Tetramer Core Facility (NTCF) and were used to stain cells for 30 minutes at room temperature (RT) before addition of other surface markers as described above.

All flow cytometry samples were acquired on a 5-laser Aurora spectral flow cytometer (Cytek Biosciences) and unmixed with Spectroflo software (Cytek). Data were analyzed using FlowJo™ v10.9 Software (BD Life Sciences).

### T cell adoptive transfers and CMV infection

Spleens from infant or adult mice were processed to generate single cell suspensions as described above. For co-transfers, a total of 5000 naïve cells containing 1:1 ratio of adult and infant OT-I T-cells were transferred into adult congenic CD45.1 host mice retro-orbitally 1 day (day -1) before infection. For single transfers, splenocytes containing 5000 OT-I CD8^+^T cells from infant or adult mice, were transferred. An equal proportion of adult:infant OT-I CD8^+^T cells was confirmed before transfer by flow cytometry. At day 0, host mice were infected i.p. with 10^4^ PFU MCMV-K181-OVA, a recombinant MCMV-K181 strain expressing full-length ovalbumin. Cells recovered from adoptive transfer experiments at indicated time-points were evaluated using flow cytometric analysis as described above.

### NK cell single cell RNA sequencing and bioinformatic data analysis

To evaluate the transcriptomic profile of NK cells prior to and following infection, splenocytes were isolated from CD45.2 infant and CD45.1 adult mice prior to infection, at day 7 post-infection, and at D15 post-infection. CD8^+^ T cells were enriched using dead cell removal kit (Miltenyi) and the mouse NK cell enrichment kit (Miltenyi) as per manufacturer protocols. For single cell profiling, cells were labeled with Biolegend CITE-Seq antibodies specific for CD45.1, and CD45.2, along with the TotalSeq-C Mouse Universal Cocktail (Biolegend) as well as a separate Hashtag (1–4) for each sample per run. CITE-Seq labeled cells were submitted to the Sulzberger Genome Center at Columbia University for encapsulation, 5’ sequencing, and alignment. Hashtags and CD45 phenotype were demultiplexed using Hashsolo; cells labeled by more than one Hashtag or exhibiting a CD45 phenotype inconsistent with the input were removed as doublets. Population assignment was performed as follows (Adult NK cells: CD45.1^+^, CD45.2^-^; Infant NK cells: CD45.1^-^, CD45.2^+^). Further doublet exclusion was performed using Scrublet. Cells were further filtered to have >1000 unique counts, >200 genes detected, and <10% of counts being mitochondrial to remove dead cells. Cells were then Leiden clustered on gene expression using Scanpy to identify NK cells (expressing *Eomes*, *Tbx21*, *Ncr1*, or *Klra8*) non-NK cells (expressing *Cd19*, *Cd5*, *Cd63,* or *Cd34*). NK cells were then again Leiden clustered with uMAP embedding. Differential gene expression was evaluated by Scanpy using Wilcoxon rank-sum. Pseudobulking was performed using decoupleR on input of cells equilibrated by both age and day post-infection.

### CD8^+^ T cell single cell RNA sequencing and bioinformatic data analysis

To evaluate the transcriptomic profile of OT-I CD8^+^ T cells prior to and following infection, cells were isolated from single cell suspensions from spleen of naïve infant and adult mice or recovered from adoptively transferred populations, as described above, at day 7 post-infection. CD8^+^ T cells were enriched using dead cell removal kit (Miltenyi) and CD8^+^ T cell enrichment kit (STEMCELL) per manufacturer protocols. A portion of the isolated cells were labeled with Biolegend CITE-Seq antibodies specific for: CD45.1, CD45.2, Thy1.1, Thy1.2, CD4, CD8a, KLRG1, and CD127 as well as a separate Hashtag (1–5) for each sample per run. CITE-Seq labeled cells were submitted to the Sulzberger Genome Center at Columbia University for encapsulation, 5’ sequencing, and alignment. Hashtags and CD45/Thy1 phenotype were demultiplexed using Hashsolo; cells labeled by more than one Hashtag or exhibiting a CD45/Thy1 phenotype inconsistent with the input were removed as doublets. Population assignment was performed as follows (Adult OT-I CD8^+^ T cells: CD45.1^-^, CD45.2^+^, Thy1.1^-^, Thy1.2^+^; Infant OT-I CD8^+^ T cells: CD45.1^-^, CD45.2^+^, Thy1.1^+^, Thy1.2^-^; and Host CD8^+^ T cells: CD45.1^+^, CD45.2^-^, Thy1.1^-^, Thy1.2^+^). Further doublet exclusion was performed using Scrublet. Cells were further filtered to have >1000 unique counts, >200 genes detected, and <10% of counts being mitochondrial to remove dead cells. Cells were then Leiden clustered on gene expression using Scanpy to identify non-CD8^+^ T cells (expressing *Cd19*, *Klrb1c*, *Cd4, Klra8*). OT-I CD8^+^ T cells were further selected by lack of expression of CD45.1 protein. OT-I CD8^+^ T cells were then again Leiden clustered with uMAP embedding. Differential gene expression was evaluated by Scanpy using Wilcoxon rank-sum. Pseudobulking was performed using decoupleR on input of cells equilibrated by both age and day post-infection.

### Defining a gene signature for human adaptive-like NK cells across tissues

We obtained single cellular indexing of transcriptome and epitope (CITE-seq) data for human innate lymphocytes including NK cells and innate lymphoid cells (ILC) from blood, lymphoid and mucosal sites using our recently published dataset (*53*). We applied score-guided Hierarchical Poisson Factorization (scHPF) independently to NK/ILC compartments from each of the six tissue types. For each tissue, we ran scHPF across a range of K values (K = 15-30 factors) to capture varying levels of transcriptional program granularity.

#### Consensus factor identification within tissues

To identify robust, reproducible factors within each tissue, we first created sparse gene loading representations by retaining only the top 1,000 genes (by score) for each factor, setting all other gene loadings to zero. We then L2-normalized these sparse gene loading vectors and computed pairwise cosine distances between all factors from different K values within each tissue. Using hierarchical clustering with average linkage on the resulting distance matrix, we identified consensus factors by optimizing the distance threshold for cutting the dendrogram. Specifically, we tested distance thresholds ranging from 0.05 to 0.9 and selected the threshold that maximized the silhouette score, ensuring biologically meaningful clustering while avoiding trivial solutions (single clusters or all singletons). We required a minimum cluster size of 3 factors to define a consensus factor, ensuring that each consensus represented a program consistently identified across multiple model parameterizations. Each consensus factor thus represents a recurring transcriptional program captured across multiple K values within that tissue.

#### Cross-tissue consensus modeling

To identify conserved transcriptional programs across tissues, we applied the same consensus procedure to the combined set of within-tissue consensus factors from all six tissues. We computed pairwise cosine distances between L2-normalized sparse gene loadings (top 1,000 genes per factor), performed hierarchical clustering with average linkage, and selected the optimal distance threshold by maximizing the silhouette score with a minimum cluster size of 3. This two-stage consensus approach enabled us to distinguish tissue-specific from tissue-agnostic NK/ILC transcriptional programs.

#### Removal of technical artifact factors

To filter out nuisance factors representing technical artifacts rather than biological signals, we performed preranked gene set enrichment analysis (GSEA) on each of the 67 consensus factors against curated gene sets for common scRNA-seq artifacts, including histone genes (HIST family) and immunoglobulin chain genes (IG genes). Factors were considered enriched for artifact signatures if they met the criteria of FDR < 0.1 and contained more than 3 genes from the artifact gene sets. Six factors meeting these criteria were excluded from downstream biological interpretation, resulting in 61 biologically informative consensus factors for subsequent analyses.

#### Association of Consensus Factors with Donor Metadata

To identify transcriptional programs associated with donor characteristics, we performed generalized linear model (GLM) analysis on the cell scores for each of the 61 biologically informative consensus factors within each tissue. For each donor-tissue combination, we first aggregated cell scores by computing the mean factor score across all cells from that donor in that tissue. We then modeled these aggregated factor scores as a function of donor-level covariates using the formula:

*y ∼ processing_site + age_groups + hcmv + sex + chemistry*

where y represents the mean cell scores for a given factor. GLMs were fit using a Gaussian family with identity link (statsmodels Python package). We applied Benjamini-Hochberg false discovery rate (FDR) correction across all tests and considered factors significantly associated with a covariate if they met the FDR < 0.05 threshold in at least one tissue.

This analysis revealed that 4 factors were significantly associated with at least one donor characteristic (FDR < 0.05). One factor associated with HCMV seronegativity showed strong enrichment for canonical myeloid markers (CD163, MARCO, MS4A6A) and myeloid-specific transcription factors (SPI1, CEBPA) rather than NK/ILC gene signatures. This likely represents technical artifacts such as doublets or misclassified myeloid contamination of the NK/ILC compartment. This factor was excluded from biological interpretation. The remaining 3 HCMV-associated factors—1 positively associated with HCMV seropositivity and 2 with HCMV seronegativity—showed gene signatures consistent with bona fide NK/ILC transcriptional programs and were carried forward for biological interpretation. The single factor enriched among the HCMV provided a basis for MMoCHi annotation of adaptive-like NK cells (*56*). Adaptive-like NK cells were then compared for differential gene expression (GEX) relative to other NK/ILC subsets as well as enrichment of protein (ADT) expression across the modeled age-coefficient.

### Human tissue acquisition and processing

We have previously detailed the processes by which we procure, process, and bank human tissue immune cells using samples from organ donors (*50*, *53*, *78*). From this banked tissue resource cells were batch thawed cells (10 separate batches) for analysis by flow cytometry (detailed above). Batches cumulatively drew from 76 distinct donors varied across age, sex, and HCMV-serostatus; including up to 7 distinct tissue sites (blood [BLD], spleen [SPL], bone marrow [BOM], hepatic lymph nodes [HLN], liver [LVR], lung lymph nodes [LLN], and lung [LNG]) from a given donor. Lymph node results did not vary by tissue location and were treated as a singular sample set for analyses [LNs]. We did not observe variance across the batches of tissue analyzed.

### Statistical Analysis

Descriptive analyses and statistical testing were performed using Prism (GraphPad ver. 10.1). Differences between two populations from separate hosts were assessed using unpaired two-tailed Student’s t-test. Differences between two populations within the same host (co-transfer set-up) were assessed by using a paired two-tailed Student’s t-test. For multiple comparisons of parameters between two groups across timepoints two-way ANOVA with Bonferroni post-test was performed. Analyses were paired for cells derived from the same individual (mouse and human). Linear regressions determined significance using Spearman correlation. Cross tissue correlations were assessed by Pearsons R. To analyze the kinetics of the adaptive-like NK cell response to infection across ages, nonlinear regression models were used. * p < 0.05.

## SUPPLEMENTARY MATERIALS

Figure S1. Flow cytometry gating strategies.

Figure S2. NK cell kinetics over age.

Figure S3. Fine-grained clustering of NK cell transcriptional states.

Figure S4. Kinetics of the CD8^+^ T cell response to MCMV infection in adult mice across sites.

Figure S5. Transcriptomic distinctions among infant and adult naive and MCMV-specific T cells in acute infection.

Figure S6. Memory and effector potential of MCMV-specific T cell populations.

Figure S7. Organ donor cohort demographics.

Figure S8. Age and tissue influence adaptive-like NK cell and CD8^+^ T_EMRA_ cell phenotypic composition.

Table S1. Adult and Infant NK cell differential gene expression at 0,7,15 DPI.

Table S2. Adult and Infant OT-I CD8 T cell differential gene expression at 0,7 DPI.

Table S3. Gene Rank of Adaptive-like NK cell scHPF signature.

Table S4. Donor Information Table

Table S5. Key resources used in this study.

## ACKNOWLEDGEMENTS

We would like to thank members of the Farber lab for their input on this project and Dr. Andrew Yates for providing critical feedback on the manuscript. Research reported in this publication was performed in the CCTI Flow Cytometry Core (NIH; S10OD030282) as well as the Sulzberger Columbia Genome Center and Single Cell Analysis Core (NIH; P30CA013696).

## Funding

This work was supported by National Institutes of Health (NIH) grants AI168634 and AI150680 awarded to D.L.F. I.J.J. was supported by T32AI148099 and D.J.C was supported by T32AI106711.

## Author Contributions

Conceptualization: IJJ, BJR, DLF. Data Curation: IJJ, BJR. Formal Analysis: IJJ, BJR, JIG, DPC. Funding acquisition: DLF. Investigation: IJJ, BJR, DLF. Methodology: IJJ, BJR, DLF. Project administration: DLF. Supervision: DLF. Visualization: IJJ, BJR. Writing – original draft: IJJ, BJR, DLF. Writing – review & editing: All authors.

## Declaration of Interests

I.J.J. has received conference travel support from Sony Biotechnology.

## Data and materials availability

Data will be made available upon reasonable request. scRNA-seq data are available at NCBI for NK cells: GEO under accession code GSE286266 and OT-I CD8+T cells: GEO under accession code GSE286265.

## Supplementary Materials for

**Figure S1.**
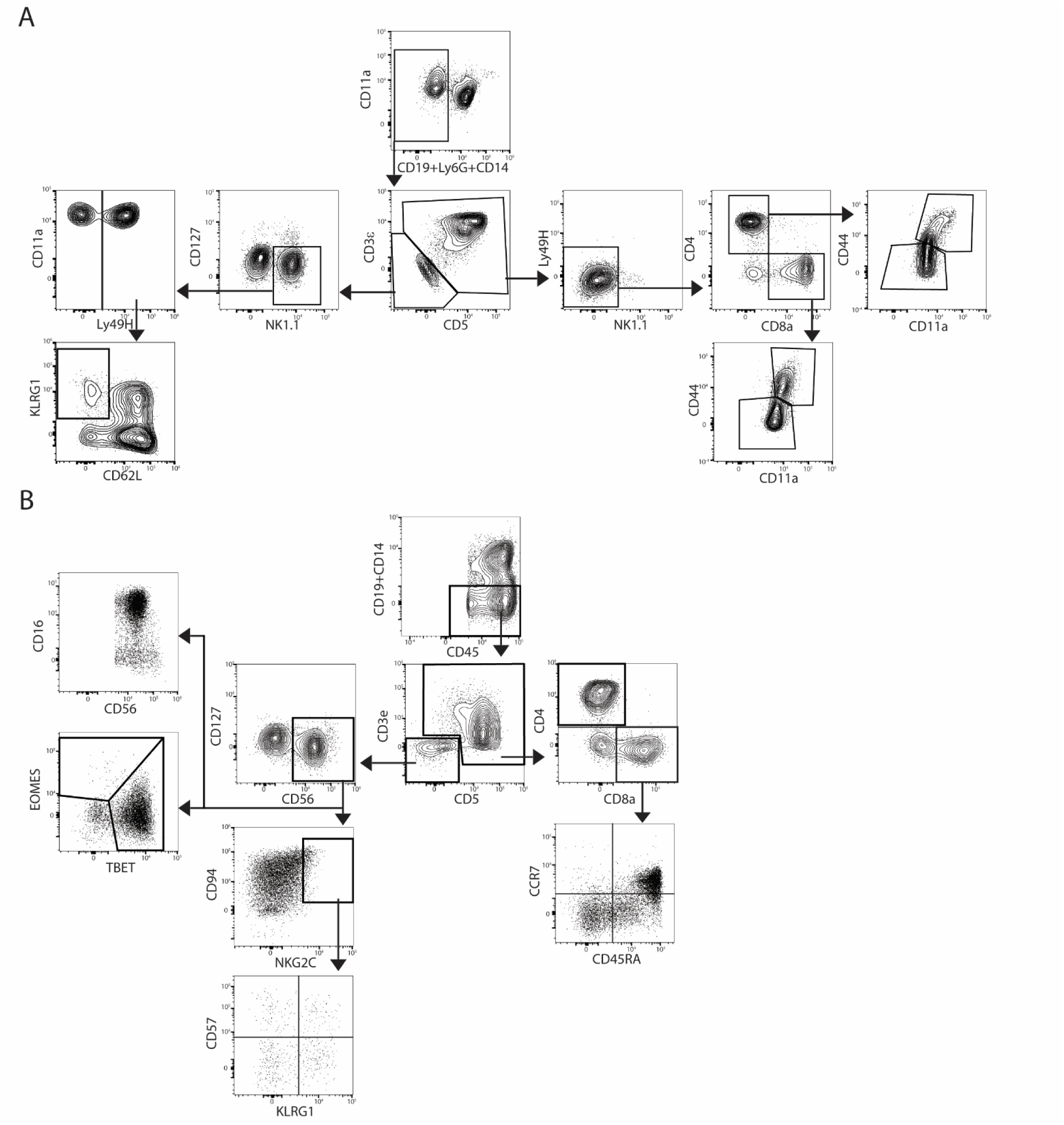
Flow cytometry gating strategies. A) Mouse flow cytometry gating strategy for NK cells (CD3^-^CD5^-^CD19^-^CD14^-^Ly6G^-^CD127^-^NK1.1^+^), identification of MCMV-reactive Ly49H^+^ NK cells and adaptive-like (Ly49H^+^KLRG1^+^CD62L^-^) NK cells, and T cells (CD3^+^ CD5^+^CD19^-^Ly6G^-^NK1.1^-^) including CD8^+^ T cells, CD4^+^ T cells, antigen-experienced (CD44^hi^CD11a^hi^), and naïve (CD44^lo^CD11a^lo^) subsets. (B) Human flow cytometry gating strategy for NK cells e^-^CD5^-^CD19^-^CD14^-^CD127^-^CD56^+^), identification of immature NK cells (CD56^bright^ CD16^-^ or EOMES^hi^ TBET^lo^), mature NK cells (CD56^dim^ CD16^+^ or EOMES^lo^ TBET^hi^), HCMV-reactive (NKG2C^+^CD94^+^) NK cells, and adaptive-like (Ly49H^+^CD94^+^[KLRG1^+^ and/or CD57^+^]) NK cells, and T cells (CD3^+^ CD5^+^CD19^-^CD14^-^) including CD8+ T cells, naive (T_N_; CD45RA^+^ CCR7^+^), central memory (T_CM_; CD45RA^-^ CCR7^+^), effector memory (T_EM_; CD45RA^-^ CCR7^-^), and effector memory expressing CD45RA (T_EMRA_; CD45RA^+^ CCR7^-^) ts.

**Figure S2.**
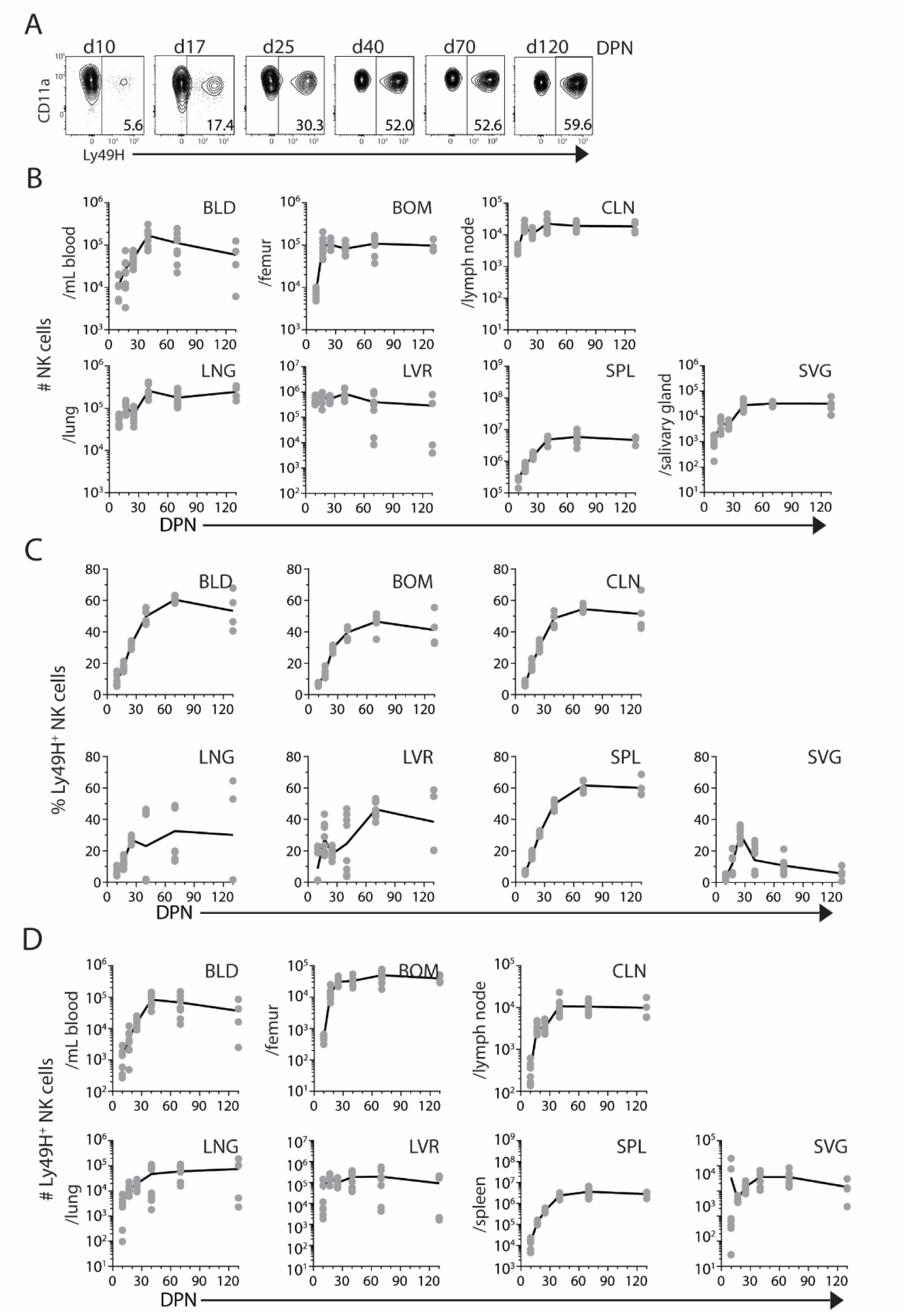
NK cell kinetics over age. (A) Representative gating for splenic Ly49H^+^ NK cells at indicated days post-natal (DPN). (B) Number of NK cells across tissues at indicated ages. (C) Frequency and (D) number of Ly49H^+^ NK cells in tissues at indicated ages. n = 8 mice per time-point per group. Data are combined from N=2 individual experiments. Tissue Abbreviations: blood (BLD), bone marrow (BOM), cervical lymph node (CLN), lung (LNG), liver (LVR), spleen (SPL), and salivary gland (SVG)

**Figure S3.**
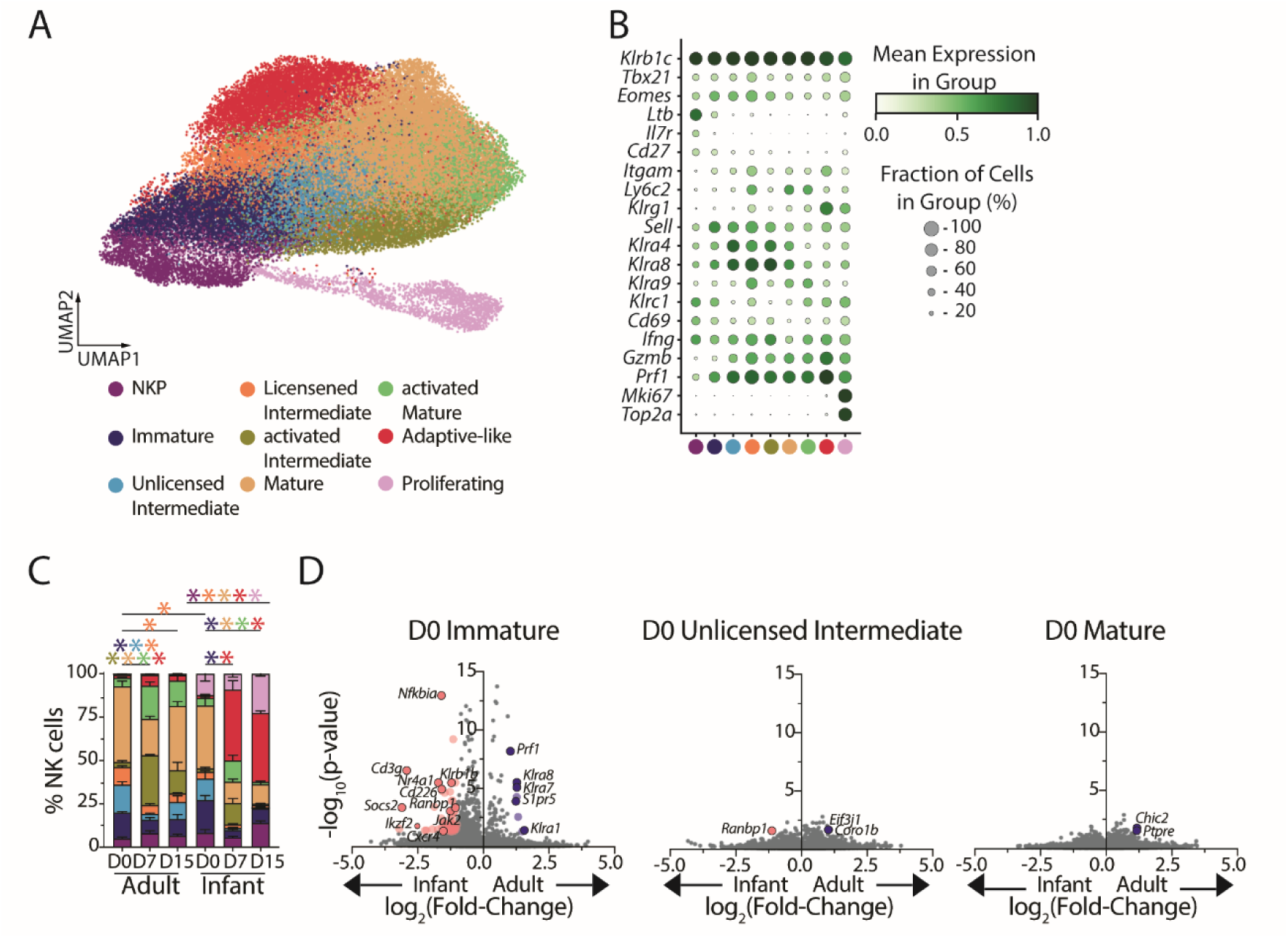
Fine-grained clustering of NK cell transcriptional states. NK cells were isolated by magnetic enrichment from the spleens of naive and adult mice prior to infection (D0) and 7 and 15 days post-MCMV infection (D7, D15) for CITE-seq profiling using the 10x genomics system. (A) Leiden clustering of adult and infant NK cells pre- and post-infection, with legend indicating putative cluster identity. (B) Dot plot displaying the relative frequency and degree of expression for indicated genes that define subsets based on Leiden clusters in panel A. (C) Proportion of each NK cell subset for adult and infant mice pre- and post-infection. n = 4 recipients per age group per timepoint; *p<0.05 as assessed by two-way ANOVA with Bonferroni multiple comparisons test; asterisk color corresponds to the population that is significantly different. (D) Volcano plots displaying differentially expressed genes (pink/blue dots; p-value <0.05, log2(fold change)>1) between adult and infant prior to infection among immature, unlicensed intermediate, and mature NK cells.

**Figure S4.**
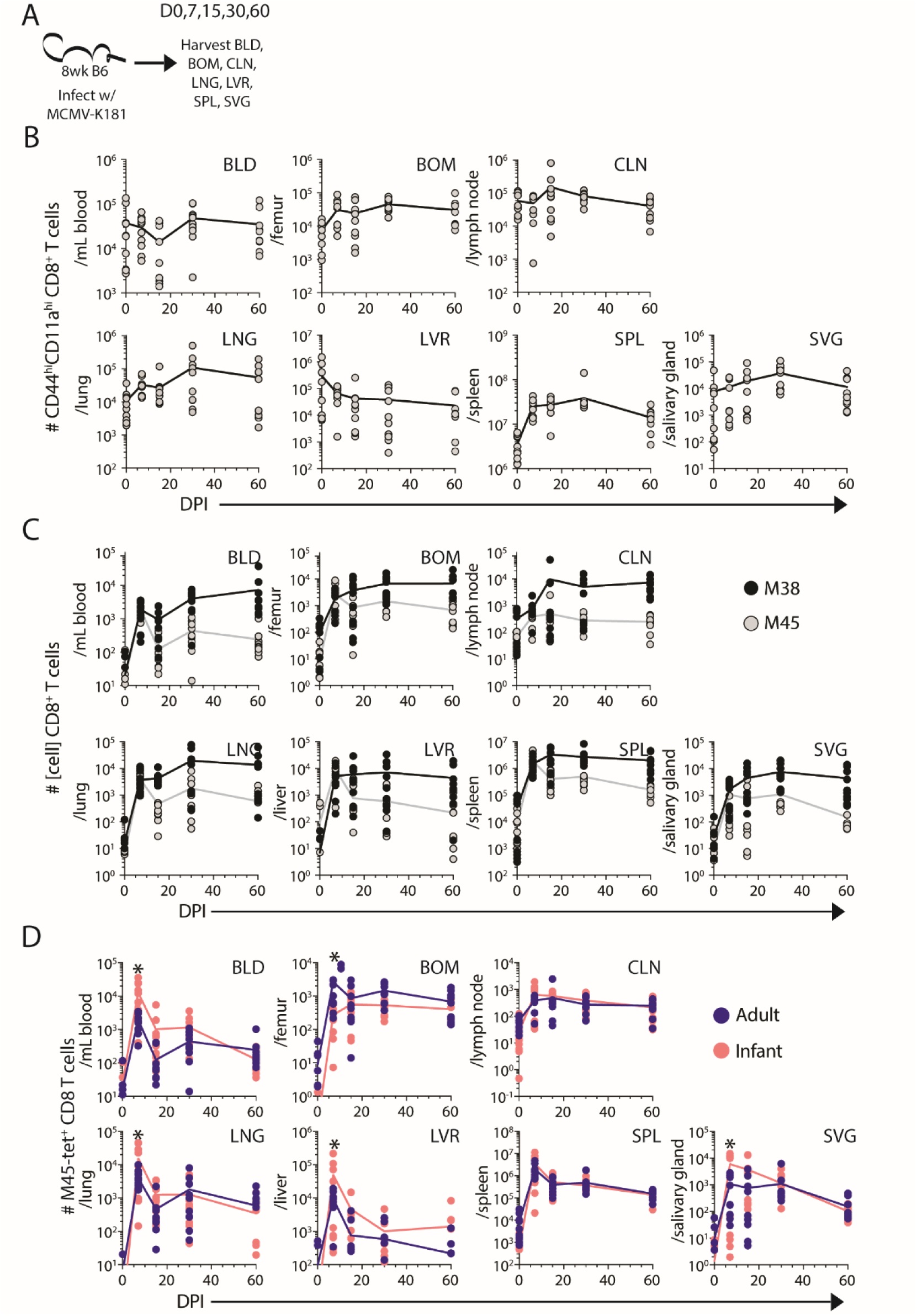
Kinetics of the CD8+ T cell response to MCMV infection in adult mice across sites. (A) Mice were infected with MCMV and blood and tissues were harvested at 0-60days post-infection. (B) Number of antigen-experienced (CD44^hi^CD11a^hi^) CD8 (circles) T cells per tissue site at indicated day post infection (DPI). (C) Number of MCMV-specific M38 (black circle) and M45 (grey circle) CD8+T cells across tissues at indicated DPI. Number of (D) M45-specific CD8^+^ T cell populations across tissues from infant (10 DPN infection; pink) and adult (70 DPN infection; blue) mice. n = 10 mice per time-point per group. * p-value < 0.05 as assessed by two-way ANOVA with Bonferroni multiple comparisons test. Data are combined from N = 2 independent experiments. Tissue Abbreviations: blood (BLD), bone marrow (BOM), cervical lymph node (CLN), lung (LNG), liver (LVR), spleen (SPL), and salivary gland (SVG)

**Figure S5.**
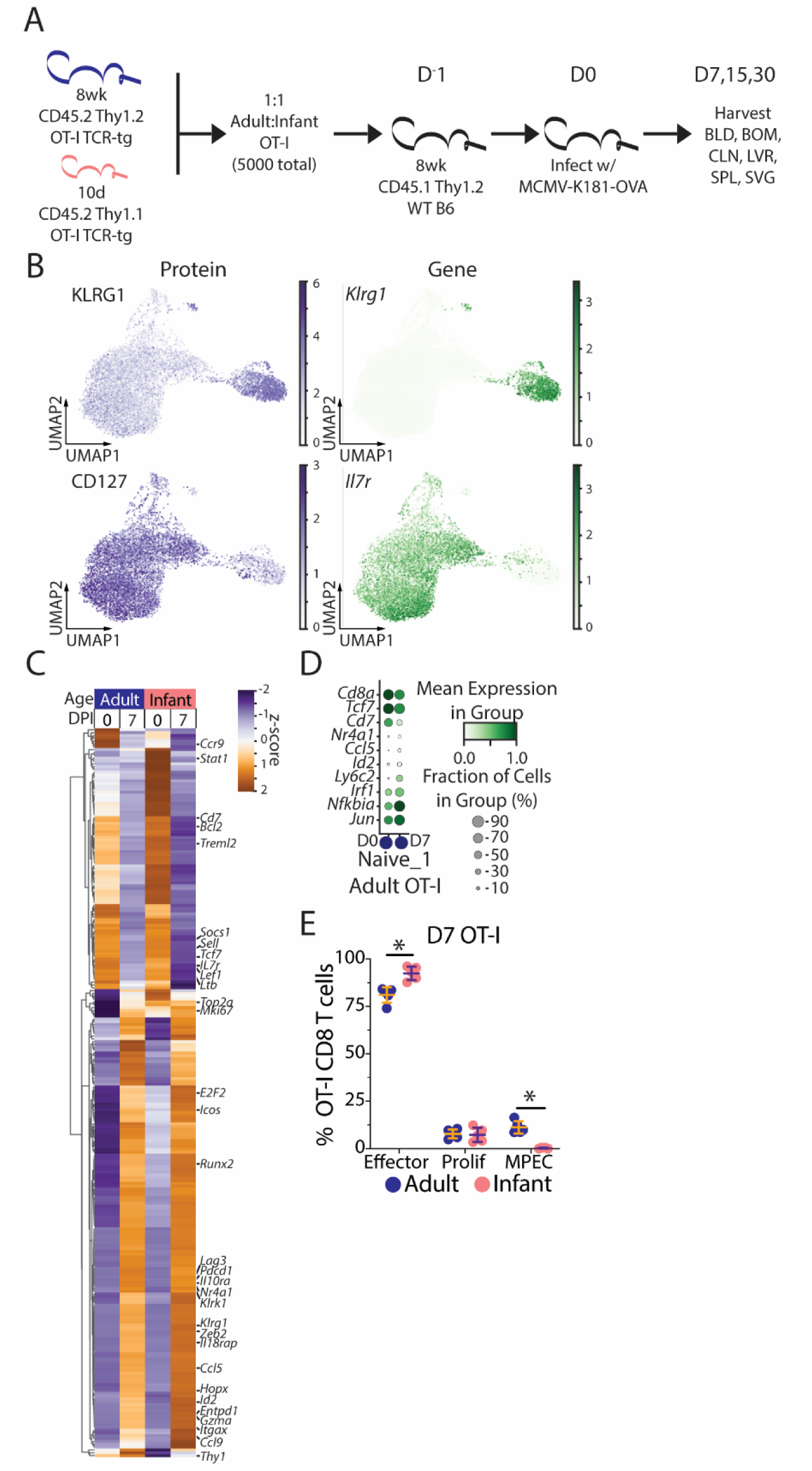
Transcriptomic distinctions among infant and adult naive and MCMV-specific T cells in acute infection. (A) Experimental schematic for co-transfer of adult and infant TCR-transgenic OT-I CD8^+^ T cells in Figure 4. (B) Expression of surface molecules bound by CITE-Seq antibodies (left) and corresponding gene expression (right) for KLRG1 (top) and CD127 (bottom). (C) Z-scored pseudobulk heatmap of the 100 most differentially expressed genes (p<0.05, log2(fold change)>1) from each group of enriched splenic OT-I CD8^+^ T cells from adult (blue) and infant (pink) mice at 0 and 7 DPI. (D) Dot plot displaying the relative frequency and degree of expression for indicated genes in adult OT-I CD8^+^ T cells in Naïve cluster 1 prior to (D0) and 7 days postinfection (D7). (E) Frequency of OT-I CD8^+^ T cells classified as effector, proliferating, or MPEC from adult (blue) and infant (pink) donors at 7 days post-infection. n = 5 recipients *p<0.05 assessed by two-way ANOVA with Bonferroni multiple comparisons test.

**Figure S6.**
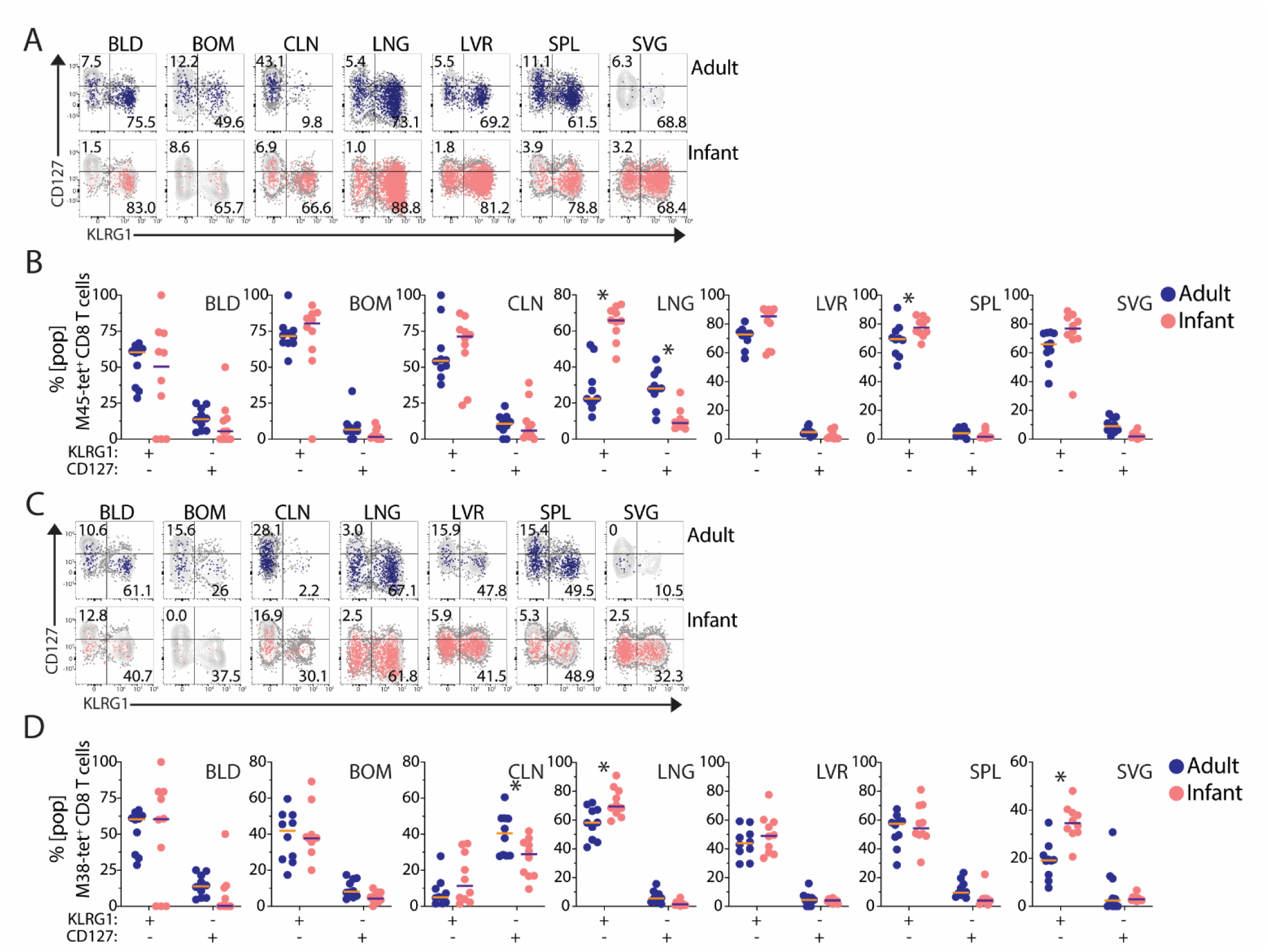
Memory and effector potential of MCMV-specific T cell populations. Frequency of CD127- KLRG1^+^ (SLEC) or CD127^+^ KLRG1- (MPEC) of adult (blue) or infant (pink) M45-tetramer^+^ cells at 7 DPI overlaid on total CD8 T cells (grey) shown in representative flow cytometry plots from indicated sites (A) and compiled data from n=10 mice per group (B). (C, D) Same as (A-B) for M38-tetramer^+^ CD8 T cells. *p < 0.05 assessed by two-way ANOVA with Bonferroni multiple comparisons test. Data are combined from N = 2 independent experiments. Tissue Abbreviations: blood (BLD), bone marrow (BOM), cervical lymph node (CLN), lung (LNG), liver (LVR), spleen (SPL), and salivary gland (SVG)

**Figure S7.**
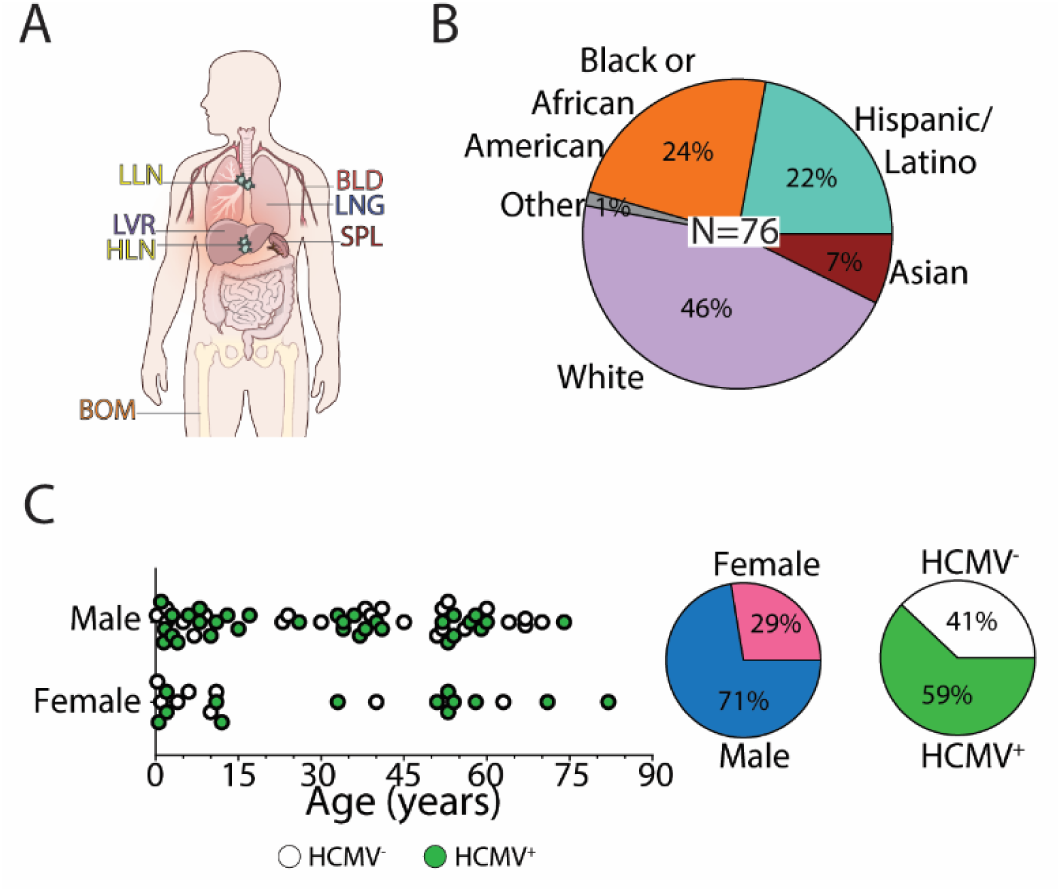
Organ donor cohort demographics. (A) Illustration of tissue sites analyzed by flow cytometry. (B) Reported ethnicities of the 76 organ donors utilized. (C) Distribution of HCMV-serostatus and Sex by Age. Tissue Abbreviations: blood (BLD), spleen (SPL), bone marrow (BOM), lymph nodes (LNs; hepatic and lung), liver (LVR), and lung (LNG)

**Figure S8.**
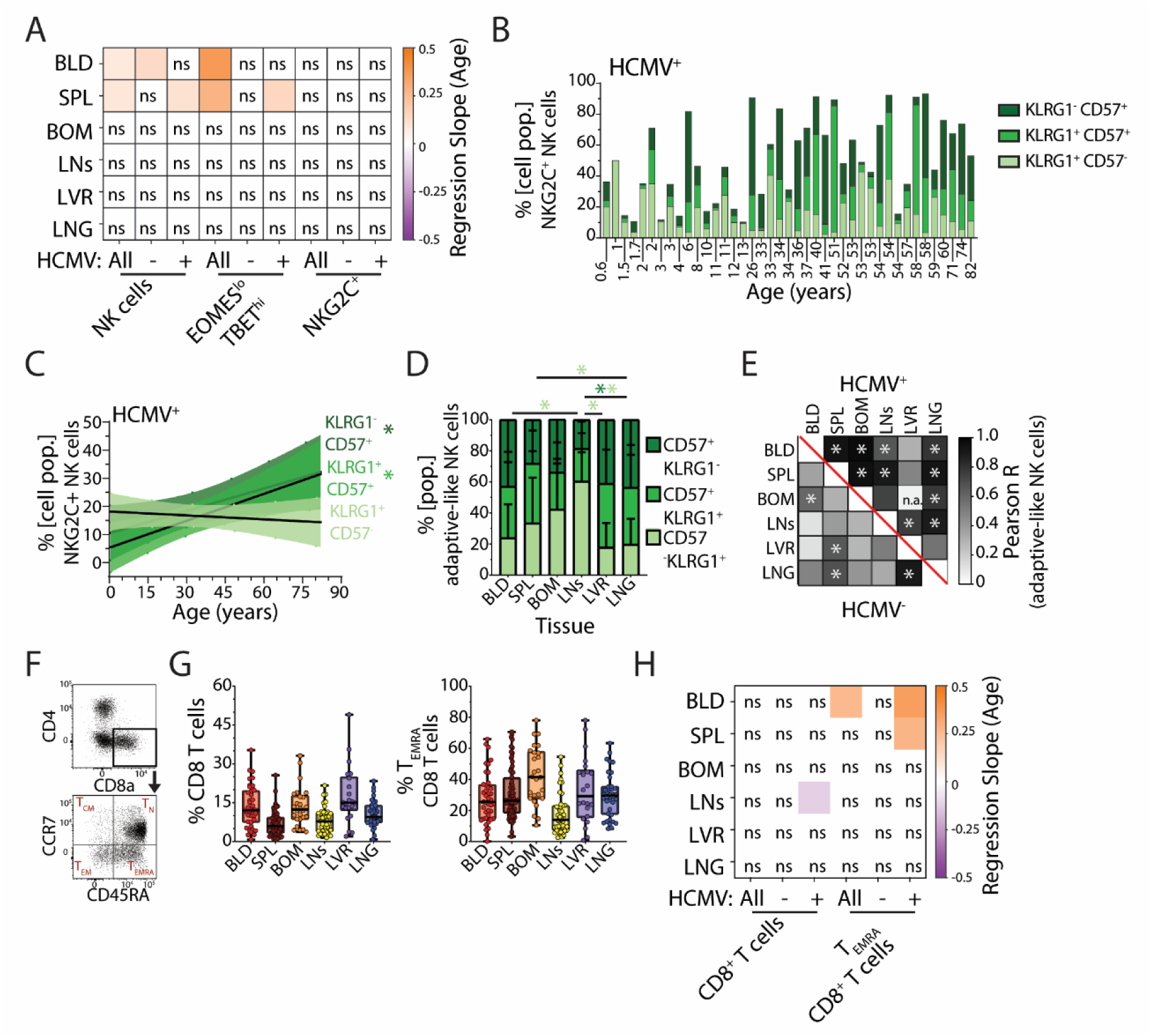
Age and tissue influence adaptive-like NK cell and CD8^+^ TEMRA cell phenotypic composition. (A) Heatmap of linear regression slopes for tissue NK cells (Figure 6I,J; Figure 7B) with significant effects of age. Orange and purple correspond to increasing and decreasing frequencies of indicated cell populations with age from all, HCMV-, and HCMV^+^ donors. (B) Composition of CD57/KLRG1 expression among NKG2C^+^ NK cells in spleen of HCMV^+^ donors. (C) Linear regression for CD57/KLRG1 populations among NKG2C^+^ NK cells by donor age in spleen of HCMV^+^ donors. Colored bands correspond to confidence intervals. *p < 0.05 (D) Frequency of KLRG1/CD57 among adaptive-like ([KLRG1^+^ or CD57^+^] NKG2C^+^) NK cells in tissues from HCMV^+^ donors. *p < 0.05 assessed by two-way ANOVA with Bonferroni multiple comparisons test and pairing tissues within donors. (E) Pearson R correlation of the frequency of adaptive-like NK cells across tissues from HCMV^+^ (top/ right) and HCMV- (bottom/ left) donors. (F) Representative gating of CD8 T cells and CD8 TEMRA cells. (G) Frequency of CD8^+^ T cells among lymphocytes (left) and TEMRA among CD8^+^ T cells (right) in indicated donor tissues. (H) Heatmap of linear regression slopes for tissue CD8 T cells (panel G; Figure 7F) with significant effects of age. Orange and purple correspond to increasing and decreasing frequencies of indicated cell populations with age from all, HCMV-, and HCMV^+^ donors. *p < 0.05. Tissue Abbreviations: blood (BLD), spleen (SPL), bone marrow (BOM), lymph nodes (LNs; hepatic and lung), liver (LVR), and lung (LNG)

**Table S1.** Adult and Infant NK cell differential gene expression at 0,7,15 DPI.

**Table S2.** Adult and Infant OT-I CD8 T cell differential gene expression at 0,7 DPI.

**Table S3.** Gene Rank of Adaptive-like NK cell scHPF signature.

**Table S4.**
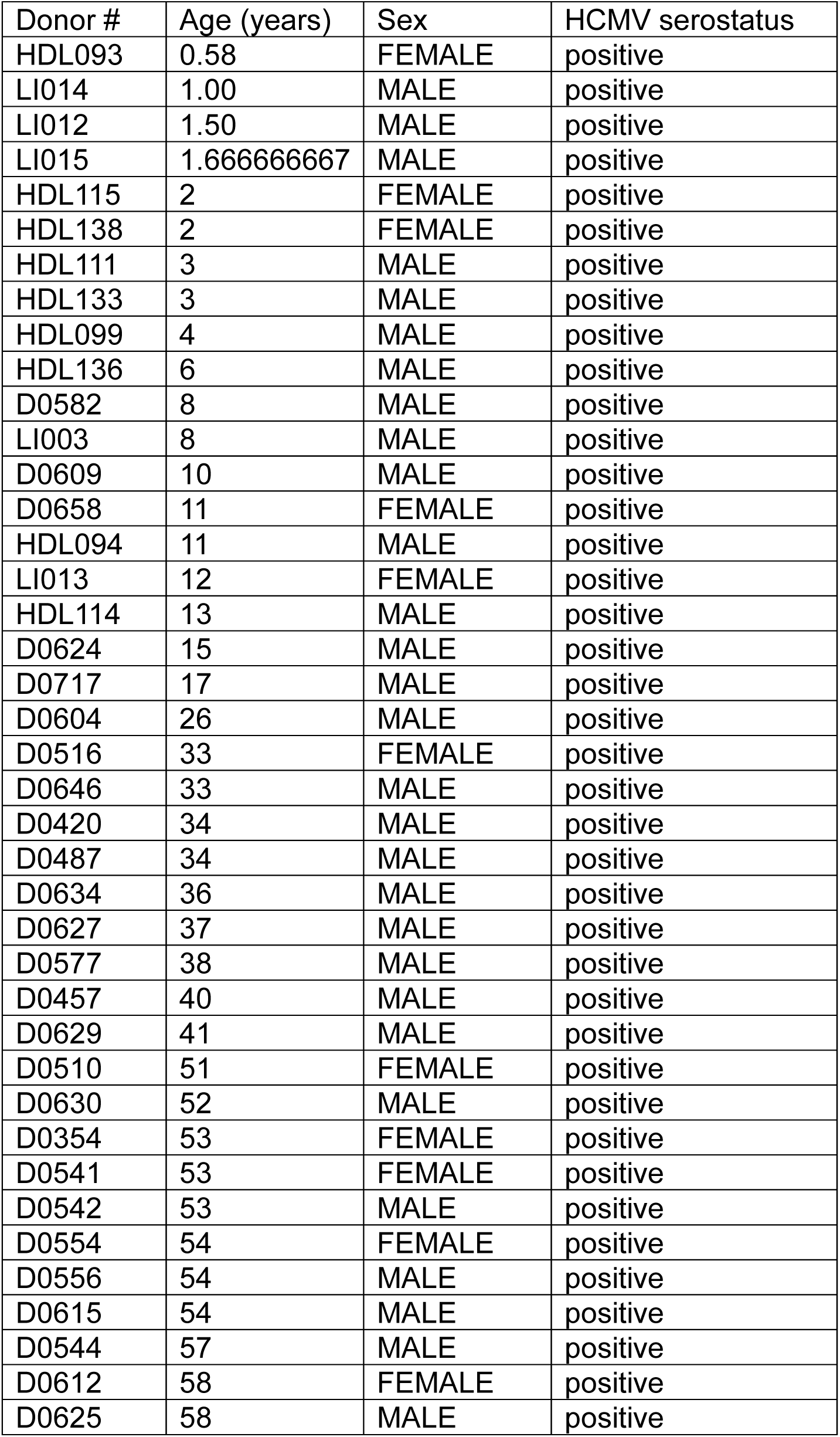
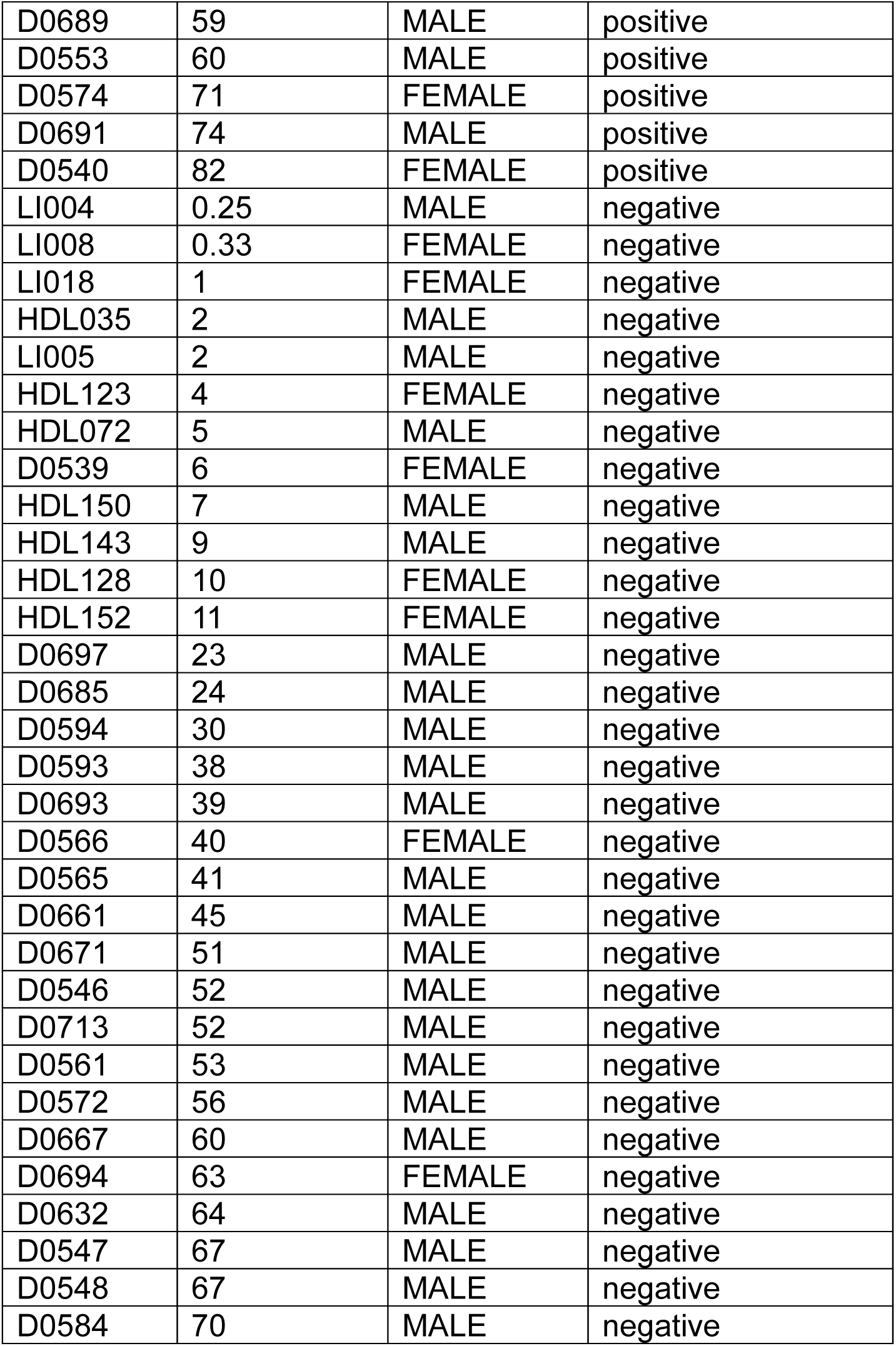
Donor Information Table.

**Table S5.** Key resources used in this study.

| Reagent or Resource | Source | Identifier |
| --- | --- | --- |
| Antibodies |  |  |
| Flow Cytometry |  |  |
| Anti-mouse CD11a BUV496 | BD | M17/4, Cat. #741071 |
| Anti-mouse CD11a PE-Cy7 | Biolegend | M17/4, Cat. #101122 |
| Anti-mouse CD127 eFlour 450 | ThermoFisher | eBioSB/199 (SB/199), Cat. #48-1273-82 |
| Anti-mouse CD14 FITC | Biolegend | Sa14-2, Cat. #123308 |
| Anti-mouse CD19 FITC | Biolegend | 6D5, Cat. #115506 |
| Anti-mouse CD3e AF700 | Biolegend | 500A2, Cat. #152316 |
| Anti-mouse CD3e PerCP-Cy5.5 | Biolegend | 500A2, Cat. #152312 |
| Anti-mouse CD4 BV605 | Biolegend | GK1.5, Cat. #100451 |
| Anti-mouse CD4 PE-Cy5 | Biolegend | RM4-5, Cat. #100513 |
| Anti-mouse CD44 BUV805 | BD | IM7, Cat. #741921 |
| Anti-mouse CD44 BV605 | Biolegend | IM7, Cat. #103047 |
| Anti-mouse CD45 APC-Fire 810 | Biolegend | 30-F11, Cat. #103174 |
| Anti-mouse CD45 BUV395 | BD | I3/2.3, Cat. #567451 |
| Anti-mouse CD45.1 APC-Fire 810 | Biolegend | A20, Cat. #110703 |
| Anti-mouse CD45.2 BV421 | Biolegend | 104, Cat. #109832 |
| Anti-mouse CD5 PerCP-Vio 700 REAfinity | Miltenyi Biotec | REA421, Cat. #130-128-069 |
| Anti-mouse CD8a BV510 | Biolegend | 53-6.7, Cat. #100752 |
| Anti-mouse CD8a BV570 | Biolegend | 53-6.7, Cat. #100740 |
| Anti-mouse CD90.1 (Thy1.1) Pacific Blue | Biolegend | OX-7, Cat. #202521 |
| Anti-rat CD90/mouse CD90.1 (Thy1.1) PE | Biolegend | OX-7, Cat. #202524 |
| Anti-mouse CD90.2 (Thy1.2) APC | Biolegend | 53-2.1, Cat. #140312 |
| Anti-mouse CD90.2 (Thy1.2) PercP-Cy5.5 | Biolegend | 30-H12, Cat. #105337 |
| Anti-mouse KLRG1 PE-Cy7 | Biolegend | 2F1/KLRG1, Cat. #138416 |
| Anti-mouse Ly49H AF647 | Biolegend | 3D10, Cat. #144710 |
| Anti-mouse Ly6G FITC | Biolegend | 1A8, Cat. #127606 |
| Anti-mouse M38 APC | NIH Tetramer Core | H-2K(b) MCMV m38 316-323<br>SSPPMFRV |
| Anti-mouse M45 PE | NIH Tetramer Core | H-2D(b) MCMV m45 985-993<br>HGIRNASFI |
| Anti-mouse NK1.1 PE-CF594 | BD | PK136, Cat. #562864 |
| Anti-mouse TCR Vα2 AF488 | Biolegend | B20.1, Cat. #127820 |
| Anti-mouse TCR Vβ5.1/2 APC | Biolegend | MR904, Cat. #139506 |
| Anti-human Eomes BUV395 | BD | X4-83, Cat. #567171 |
| Anti-human CD94 BUV496 | BD | HP-3D9, Cat. #750232 |
| Anti-human CD14 BUV615 | BD | M5E2, Cat. #751150 |
| Anti-human CD19 BUV615 | BD | SJ254C1, Cat. #612989 |
| Anti-human NKG2C BUV805 | BD | 134591, Cat. #749683 |
| Anti-human Tbet BV421 | Biolegend | 4B10, Cat. #644816 |
| Anti-human CD5 Pacific Blue | Biolegend | UCHT2, Cat. # 300624 |
| Anti-human CD57 BV510 | Biolegend | QA17A04, Cat. #393314 |
| Anti-human CD8a BV570 | Biolegend | RPA-T8, Cat. #301038 |
| Anti-human CD4 BV605 | Biolegend | RPA-T4, Cat. #300556 |
| Anti-human CD45RA BV711 | Biolegend | HI100, Cat. #304138 |
| Anti-human CD127 AF532 | Thermo-Fisher | EBioRDR5, Cat. #58-1278-42 |
| Anti-human CD3 NovaFluor Blue 610-70S | Thermo-Fisher | UCHT2, Cat. #H002T03B06-A |
| Anti-human CD56 PE-CF594 | BD | NCAM16.2, Cat. #564849 |
| Anti-human CD16 PE-Cy5 | Biolegend | 3G8, Cat. #302010 |
| Anti-human KLRG1 PE-Cy7 | Biolegend | SA231A2, Cat. #367720 |
| Anti-human CCR7 APC-R700 | BD | 3D12, Cat. #750232 |
| Anti-human CD45 APC-Fire810 | Biolegend | HI30, Cat. #304076 |
| Enrichment kits |  |  |
| Dead cell removal kit | Miltenyi Biotec | Cat. #130-090-101 |
| NK cell Isolation Kit, mouse | Miltenyi Biotec | Cat. #130-115-818 |
| EasySep™ Mouse CD8<sup>+</sup> T cell Isolation Kit | STEMCELL Technologies | Cat. #19853 |
| CITE-Seq Antibodies |  |  |
| TotalSeq-C0002 anti-mouse CD8a | Biolegend | 53-6.7, Cat. #100785 |
| TotalSeq-C0157 anti-mouse CD45.2 | Biolegend | 104, Cat. #109855 |
| TotalSeq-C0178 anti-mouse CD45.1 | Biolegend | A20, Cat. #11757 |
| TotalSeq-C0001 anti-mouse CD4 | Biolegend | RM4-5, Cat. #100571 |
| TotalSeq-C0075 anti-mouse CD90.2 | Biolegend | 30-H12, Cat. #105353 |
| TotalSeq-C0380 anti-mouse CD90.1 | Biolegend | OX-7, Cat. #202551 |
| TotalSeq-C0198 anti-mouse CD127 | Biolegend | A7R34, Cat. #135047 |
| TotalSeq-C0002 anti-mouse/human KLRG1 | Biolegend | 2F1/KLRG1, Cat. #138433 |
| TotalSeq-C0301 anti-mouse Hashtag 1 | Biolegend | Cat. #155861 |
| TotalSeq-C0302 anti-mouse Hashtag 2 | Biolegend | Cat. #155863 |
| TotalSeq-C0303 anti-mouse Hashtag 3 | Biolegend | Cat. #155865 |
| TotalSeq-C0304 anti-mouse Hashtag 4 | Biolegend | Cat. #155867 |
| TotalSeq-C0305 anti-mouse Hashtag 5 | Biolegend | Cat. #155869 |
| TotalSeq-C Mouse Universal Cocktail, V1.0 | Biolegend | Cat. #199903 |
| Cell Depletions |  |  |
| InVivoMAb Anti-mouse CD3e depleting antibody 145-2C11 | BioXcell | Cat. #BE0001-1 |
| InVivoMAb Anti-mouse NK1.1 depleting antibody PK136 | BioXcell | Cat. #BE0036 |
| InVivoMAb Armenian Hamster IgG Isotype PIP | BioXcell | Cat. #BE0260 |
| InVivoMAb Mouse IgG2a Isotype C1.18.4 | BioXcell | Cat. #BE0085 |
| Chemicals and Reagents |  |  |
| ACK Lysing buffer | Gibco | Ref. #A10492-01 |
| Buffer ATL | Qiagen | Cat. #939011 |
| Collagenase | Millipore Sigma | Cat. #11088882001 |
| DNase | Millipore Sigma | Cat. #DN25-5G |
| DNeasy Blood & Tissue Kit (250) | Qiagen | Cat. #69506 |
| DPBS | Corning | Cat. #20-030-CV |
| EDTA | Corning | Cat. #46-034-CI |
| Fetal Bovine Serum | GeminiBio | Cat. #100-106 |
| Fixable Viability Dye Zombie NIR | Biolegend | Cat. #423106 |
| Fixation Buffer | Invitrogen | Cat. #00-5223-56 |
| Foxp3 / Transcription Factor Fix/Perm Diluent (1X) | FisherScientific | Ref. #TNB-1022-L160 |
| IMDM | Gibco | Ref#122440-053 |
| Ionomycin | Sigma | Cat#I9657 |
| Penicillin/Streptomycin/L-glutamine | GeminiBio | Cat#c400-110 |
| Proteinase K | Qiagen | Cat. #19133 |
| RPMI 1640 | Corning | Cat. #10-040-CM |
| Solution 13 AO/DAPI Staining Reagent | Chemometec | Cat. # 910-3013 |
| SsoAdvanced Universal SYBR Green Supermix® | BioRad | Cat. #1725274 |
| Tonbo Flow Cytometry Perm Buffer (10X) | FisherScientific | Ref. #TNB-1213-L150 |
| Tonbo Foxp3 / Transcription Factor Fix/Perm Concentrate (4X) | FisherScientific | Ref. #TNB-1020-L050 |
| TruStain FcX PLUS (anti-mouse CD16/32) | Biolegend | Cat. #156604 |
| Software and Algorithms |  |  |
| FlowJo™ v 10.9 software | BD | https://www.flowjo.com |
| SpectroFlo® 3.1.0 software | CytekBio | https://www.cytekbio.com/pages/spectro-flo |
| Prism v 10.6 software | Graphpad | https://www.graphpad.com |
| BR.io cloud platform | BioRad | https://www.br.io |

